# Interface-acting nucleotide controls polymerization dynamics at microtubule plus- and minus-ends

**DOI:** 10.1101/2023.05.03.539131

**Authors:** Lauren A McCormick, Joseph M Cleary, William O Hancock, Luke M Rice

**Author notes:** Equal contributions (Lauren A McCormick, Joseph M Cleary). Equal contributions.

## Abstract

GTP-tubulin is preferentially incorporated at growing microtubule ends, but the biochemical mechanism by which the bound nucleotide regulates the strength of tubulin:tubulin interactions is debated. The ‘self-acting’ (cis) model posits that the nucleotide (GTP or GDP) bound to a particular tubulin dictates how strongly that tubulin interacts, whereas the ‘interface-acting’ (trans) model posits that the nucleotide at the interface of two tubulin dimers is the determinant. We identified a testable difference between these mechanisms using mixed nucleotide simulations of microtubule elongation: with self-acting nucleotide, plus- and minus-end growth rates decreased in the same proportion to the amount of GDP-tubulin, whereas with interface-acting nucleotide, plus-end growth rates decreased disproportionately. We then experimentally measured plus- and minus-end elongation rates in mixed nucleotides and observed a disproportionate effect of GDP-tubulin on plus-end growth rates. Simulations of microtubule growth were consistent with GDP-tubulin binding at and ‘poisoning’ plus-ends but not at minus-ends. Quantitative agreement between simulations and experiments required nucleotide exchange at terminal plus-end subunits to mitigate the poisoning effect of GDP-tubulin there. Our results indicate that the interfacial nucleotide determines tubulin:tubulin interaction strength, thereby settling a longstanding debate over the effect of nucleotide state on microtubule dynamics.

## Introduction

Microtubules are dynamic polymers of αβ-tubulin that support motor-based transport of cargo through the cytoplasm and orchestrate the movement of chromosomes in dividing cells (Akhmanova & Kapitein, 2022; Barlan & Gelfand, 2017; Cleary & Hancock, 2021; Gudimchuk & McIntosh, 2021; Prosser & Pelletier, 2017). Microtubules grow by the addition of GTP-bound tubulin to the polymer ends. Once incorporated into the microtubule lattice, tubulins hydrolyze their bound GTP. The change in nucleotide state triggers conformational changes that weaken interactions between neighboring tubulins and ultimately results in catastrophe, the switch from growth to shrinkage (Bowne-Anderson et al., 2013; Gudimchuk & McIntosh, 2021; LaFrance et al., 2022; Manka & Moores, 2018; Roostalu et al., 2020; Seetapun et al., 2012; Zanic et al., 2013). Defining the connection between nucleotide state and tubulin:microtubule binding kinetics is crucial for understanding how microtubules grow and how they transition to catastrophe. However, the mechanism by which nucleotide controls the strength of tubulin:tubulin interactions remains debated.

An early model explained the nucleotide-dependence of microtubule stability by positing that nucleotide state determines the conformation of tubulin: GTP-tubulin would form strong lattice contacts because GTP favors a ‘straight’ conformation compatible with the microtubule lattice, and GDP-tubulin would form weak lattice contacts because GDP favors a ‘curved’ conformation incompatible with the microtubule lattice (Drechsel & Kirschner, 1994; Howard & Timasheff, 1986; Melki et al., 1989; Nicholson et al., 1999; Shearwin et al., 1994; Tran et al., 1997; Wang & Nogales, 2005). By assuming that nucleotide controls the conformation of the tubulin to which it is bound, this model embodied a ‘cis-acting’ view of nucleotide action. However, subsequent work demonstrated that both GTP- and GDP-tubulin adopt the same curved conformation (Nawrotek et al., 2011; Pecqueur et al., 2012; Rice et al., 2008), which contradicted a core assumption of the cis-acting model. These structural findings led to the proposal of a ‘trans-acting’ mechanism in which the nucleotide bound to one tubulin controls the strength of its interactions with the next tubulin through direct contacts and/or by causing loop movements that lead to better polymerization contacts (Ayaz et al., 2012; Buey et al., 2006; Nawrotek et al., 2011; Piedra et al., 2016; Rice et al., 2008). The ‘trans’ mechanism is supported by the knowledge that the nucleotide binding site on β-tubulin forms part of the polymerization interface with the α-tubulin from the next subunit in the protofilament, and it is also consistent with the largest nucleotide-dependent conformational changes in the microtubule occurring in α-tubulin adjacent to the β-tubulin-bound nucleotide (Alushin et al., 2014; Manka & Moores, 2018; Zhang et al., 2015). However, the field has still not reached a consensus on the mechanism of nucleotide action (Brouhard, 2015; Brouhard & Rice, 2018; Brun et al., 2009; Luo et al., 2023; Margolin et al., 2012; Schmidt & Kierfeld, 2021; Stewman et al., 2020; VanBuren et al., 2005; VanBuren et al., 2002; Zakharov et al., 2015) and this persistent ambiguity about how nucleotide state influences tubulin:tubulin interactions limits our understanding of microtubule dynamics.

Microtubule plus- and minus-ends are structurally distinct: plus-ends present a β-tubulin polymerization interface that contains the exchangeable nucleotide, whereas minus-ends present an α-tubulin polymerization interface that does not expose a nucleotide. Debate over cis- and trans-acting mechanisms (reviewed in (Gudimchuk & McIntosh, 2021)) has persisted in part because most studies have focused solely on the plus-end, where two nucleotides – one bound to the terminal tubulin (the cis nucleotide), and one at the interface between the terminal tubulin and the next subunit in the microtubule lattice (the trans nucleotide) – could in principle be dictating the strength of lattice contacts. At the minus-end, by contrast, the exchangeable nucleotide of the terminal tubulin is already buried in the microtubule lattice. We reasoned that this fundamental difference between the plus- and minus-ends might provide a new way to test the conflicting mechanisms of nucleotide action. A few studies have compared plus- and minus-end dynamics (Strothman et al., 2019; Tanaka-Takiguchi et al., 1998; Walker et al., 1988), but only one of them sought to manipulate nucleotide state in a controlled manner (Tanaka-Takiguchi et al., 1998). For generality in considering both ends, we will hereafter refer to the trans mechanism as ‘interface-acting’, and the cis mechanism as ‘self-acting’.

The goal of the present study was to determine whether self-acting or interface-acting mechanisms of nucleotide action govern the strength of tubulin:tubulin contacts in the microtubule. Our approach used simulations and experiments to compare how plus- and minus-end elongation are affected by GDP-tubulin. We first simulated microtubule elongation in mixed nucleotide states using models that implemented self- or interface-acting mechanisms. These simulations revealed a striking difference between the two mechanisms of nucleotide action: in the self-acting model, GDP-tubulin inhibited plus- and minus-ended growth to the same extent, but in the interface-acting model, GDP-tubulin disproportionately inhibited plus-ended growth. This observation was consistent with an earlier study that showed a selective suppression of plus-end elongation when GDP was included in the reaction mixture (Tanaka-Takiguchi et al., 1998). However, the reaction conditions in that earlier study did not suppress GTPase activity, and the consequent high frequency of plus-end catastrophe prevented an examination of how GDP affected microtubule growth rates. We tested our predictions experimentally using ‘mixed nucleotide’ assays (containing both slowly-hydrolyzable GMPCPP and GDP) to prevent catastrophe, allowing us to directly compare the relative effects of GDP-tubulin on plus- and minus-end growth rates. We found that plus-end growth was disproportionately affected by GDP-tubulin, providing strong new evidence in support of the interface-acting mechanism. Further simulations revealed that nucleotide exchange can modulate the magnitude of plus-end poisoning by GDP-tubulin (Cleary et al., 2022; Piedra et al., 2016; Vandecandelaere et al., 1995). By ruling out a self-acting (cis-acting) mechanism of nucleotide action, our findings provide new evidence that resolves a longstanding debate about how the bound nucleotide governs the tubulin:tubulin interactions that dictate microtubule growth.

## Results

### Self- and interface-acting mechanisms of nucleotide action predict different effects of GDP-tubulin on plus- and minus-end growth

The self- and interface-acting mechanisms for how nucleotide dictates the strength of tubulin:tubulin interactions are illustrated in Figure 1A: the self-acting mechanism posits that the nucleotide bound to β-tubulin (GTP or GDP) controls how tightly *that* tubulin interacts with the lattice, whereas the interface-acting mechanism posits that the nucleotide at the *interface between* tubulin dimers controls how tightly they interact. At the plus-end, the two mechanisms can lead to different outcomes because there are two nucleotides involved – one bound to the terminal tubulin and one at the interface below (Figure 1A, top panels). At the minus-end, however, the two mechanisms are indistinguishable because there is only one nucleotide involved: the nucleotide bound to the terminal subunit is also the nucleotide at the interface with the lattice (Figure 1A, bottom panels). Using kinetic simulations of microtubule elongation, we sought to identify a testable difference between the self- and interface-acting mechanisms. We first expanded our model (Ayaz et al., 2014; Cleary et al., 2022; Kim & Rice, 2019; Piedra et al., 2016) to simulate both plus- and minus-end elongation and to include multiple nucleotide states for unpolymerized tubulin (summarized in Figure 1 – figure supplement 1). We then used the model to predict how GDP-tubulin might affect plus- and minus-end elongation with either the self- or interface-acting mechanisms of nucleotide action.

**Figure 1.**
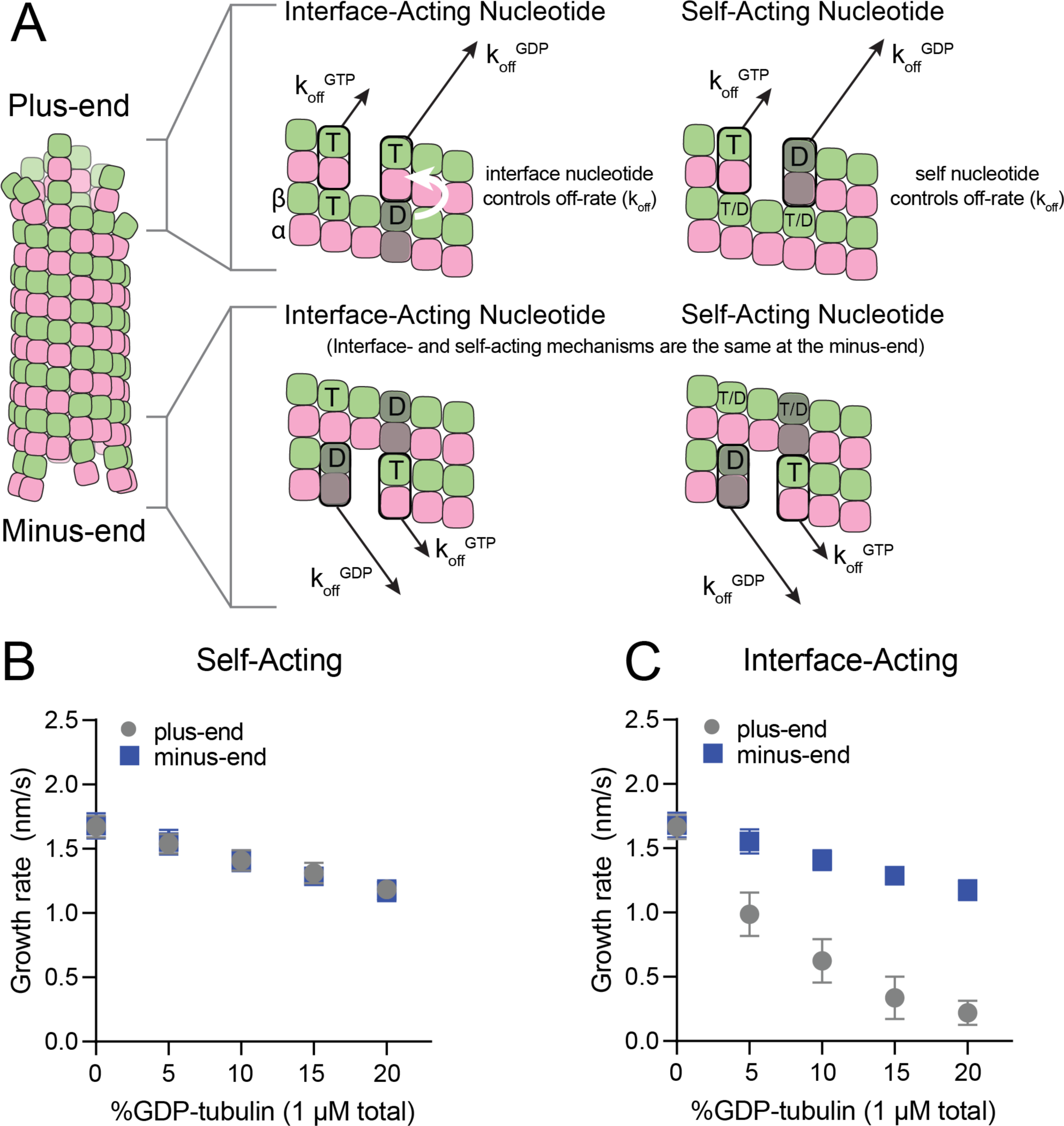
Mechanisms of nucleotide action and simulations of plus- and minus-ends. **(A)** Cartoon showing self- (cis) or interface-acting (trans) nucleotide mechanisms. In an interface-acting mechanism, the nucleotide at the interface of two tubulin dimers controls their interaction affinity, shown by a white arrow. In a self-acting mechanism, the nucleotide bound to the terminal tubulin controls how tightly that tubulin interacts with the lattice. At the plus-end, the two mechanisms can lead to different outcomes because there are two nucleotides involved – one bound to the terminal β-tubulin, and one at the interface between the terminal tubulin and the microtubule lattice. At the minus-end, however, self-acting and interface-acting mechanisms are equivalent because the incoming nucleotide becomes the interfacial nucleotide. T = GTP, T/D = GTP or GDP, D = GDP. **(B and C)** Simulated growth rates of GTP microtubule plus- and minus-ends, using arbitrarily chosen parameters that support elongation in the chosen concentration range. **(B)** In a self-acting mechanism, both plus-end (circles) and minus-end (squares) growth rates are predicted to decrease linearly with the amount of GDP-tubulin. **(C)** In an interface-acting mechanism, plus-end (circles) growth rates are predicted to be disproportionately impacted by GDP-tubulin relative to minus-end growth rates. Error bars are standard deviation (n = 50 per condition) and if not visible, are obscured by the symbols. Simulation parameters are: k_on_: 1.0 µM^-1^ s^-1^, K_D_^long^ = 100 µM, K_D_^corner^ = 100 nM, K_D_^long,GDP^ = 300 mM. The predicted difference between mechanisms at the plus-end is robust across different choices for K_D_^long^, K_D_^corner^, and the GDP weakening effect (Figure 1 Supplement 3). Note that because the two mechanisms are equivalent at the minus-end, interface-acting simulations for the minus-end use the same simulation results as the self-acting simulations. The total [tubulin] is constant, thus minus-end growth rates decrease in proportion to the decrease in the concentration of GMPCPP-tubulin.

We performed ‘mixed nucleotide’ simulations of plus- and minus-end growth at 1 µM total tubulin with varying fractions of GDP-tubulin (0-20%). For simplicity, simulations used arbitrarily chosen parameters that supported elongation in the chosen concentration regime (Table 1). To provide the simplest possible biochemical setting and to set the stage for experiments described below, simulations did not attempt to explicitly model different conformations of αβ-tubulin (see Discussion), and also ignored GTP hydrolysis. For both self-acting and interface-acting nucleotide, simulated minus-end growth rates decreased identically and in linear proportion to the amount of GDP-tubulin in the simulation (Figure 1B). However, simulated plus-end growth rates decreased much more for interface-acting nucleotide than for self-acting nucleotide (Figure 1C). Similar results were obtained for alternative parameter choices (Figure 1 Supplement 2 and 3). Based on these robust end-specific differences from simulations, comparative measurements of how GDP-tubulin affects plus- and minus-end elongation should provide a new way to test the self- or interface-acting mechanisms of nucleotide action.

**Table 1.** Simulation Parameters for Figures 1 – 5. Dashed lines denote figure panels where either plus- end or minus-end simulations were not performed; these panels focused on simulations of only one end.

| | $k_{on}^{plus}$<br>($\mu M^{-1} s^{-1}$) | $k_{on}^{minus}$<br>($\mu M^{-1} s^{-1}$) | $K_D^{long}$<br>($\mu M$) | $K_D^{corner}$<br>($\mu M$) | $K_D^{long} GDP$<br>($\mu M$) | $K_D^{corner} GDP$<br>($\mu M$) | Tubulin<br>($\mu M$) |
| --- | --- | --- | --- | --- | --- | --- | --- |
| **Fig. 1** | 1.0 | 1.0 | 100 | 0.1 | $3 \times 10^5$ | 300 | 1 |
| **Fig. 4A** | 0.74 | 0.31 | 86 | 0.025 | $3 \times 10^5$ | 87 | 1.25 |
| **Fig. 4B** | 0.74 | --- | 86 | 0.025 | $3 \times 10^5$ | 87 | 1.25 |
| **Fig. 4C** | --- | 0.31 | 86 | 0.025 | $3 \times 10^5$ | 87 | 1.25 |
| **Fig. 5B** | 0.74 | --- | 86 | 0.025 | $3 \times 10^5$ | 87 | 1.25 |

### Mixed nucleotide experiments reveal different effects of GDP-tubulin on plus- and minus-ends

To establish a baseline for measurements with mixed nucleotides, we first used interference reflection microscopy (IRM) to measure plus-and minus-end growth rates in 1 mM GMPCPP (a slowly hydrolyzable GTP analog) at multiple concentrations of bovine brain tubulin. Growth rates displayed the expected linear dependence on tubulin concentration (Figure 2A). Both ends showed the same apparent critical concentration (C_capp_) of 50 nM, but plus-end growth showed a roughly two-fold higher apparent on-rate constant (k_onapp_) than minus-end growth, 3 µM^-1^s^-1^MT^-1^ and 1.5 µM^-1^s^-1^MT^-1^ respectively (Figure 2B).

**Figure 2.**
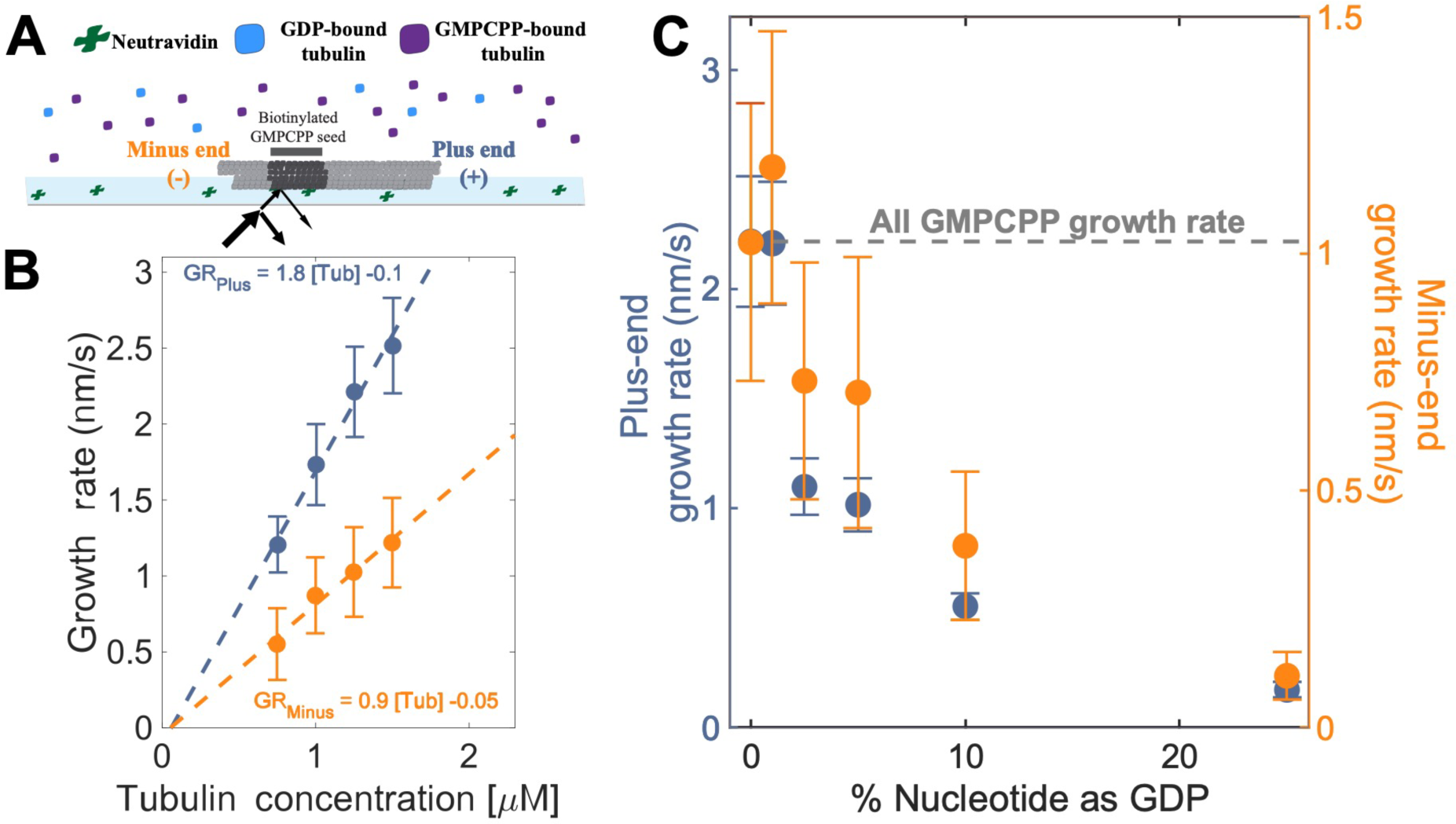
Microtubule plus- and minus-end growth both decrease in the presence of GDP-tubulin. **(A)** Schematic of the *in vitro* assay, in which biotinylated GMPCPP microtubule seeds are attached to a neutravidin coated-cover slip, and microtubule assembly in the presence of tubulin bound to either GDP- or GMPCPP is monitored using Interference Reflection Microscopy (IRM). **(B)** Growth rates of microtubule plus- and minus-ends in GMPCPP as a function of tubulin concentration. (n= 64-125 for the plus-end and n= 39-95 for the minus-end). The error bars denote standard deviation. **(C)** Plus- (left y-axis) and minus- end (right y-axis) growth rates at 1.25 µM tubulin in mixtures of GDP and GMPCPP containing 1 mM total nucleotide. The gray line denotes the “all GMPCPP” growth rates of the two ends. Growth rates are normalized by the growth rate at 1.25 µM tubulin in 1 mM GMPCPP (n = 66 - 121 for the plus-end and n = 44 - 94 for the minus-end). The error bars denote standard deviation. Using a two-sided t-test with unequal variance, differences in the mean normalized growth rates at plus- and minus-ends were statistically significant with p < 0.001 for all nucleotide mixtures except 0% GDP.

As a way to test the predictions from our simulations (Figure 1BC), we next measured the growth rates of microtubule plus- and minus-ends at a constant concentration of tubulin (1.25 µM) but using different ratios of GDP and GMPCPP (1 mM total nucleotide concentration). Growth rates at both ends decreased substantially in mixtures containing as little as 2.5% GDP (25 µM GDP and 975 µM GMPCPP) (Figure 2C), but plus-end growth rates decreased to a greater degree than minus-end growth rates. For instance, at 25 µM GDP, plus-end growth rates fell ∼50% (from 2.2 nm/s to 1.1 nm/s) relative to “all GMPCPP” growth rates, whereas minus-end growth rates only fell ∼30% (from 1 nm/s to 0.7 nm/s). This ∼1.5-fold stronger inhibition by GDP of plus-end growth rates held across multiple nucleotide mixing ratios (Fig. 2C). Importantly, the larger decrease in the growth rate at the plus-end agrees with the predictions made by the interface-acting mechanism (Figure 1A) compared to the self-acting mechanism.

### Plus-end growth is super-stoichiometrically suppressed by GDP-tubulin

Tubulin binds different nucleotides with different affinities (Aldaz et al., 2005; Chakrabarti et al., 2000; Correia et al., 1987; Fishback & Yarbrough, 1984; Hyman et al., 1992; Mejillano & Himes, 1991; Monasterio & Timasheff, 1987; Zeeberg & Caplow, 1979), so the ratio of GDP and GMPCPP in a reaction does not directly translate to the fractions of GDP- and GMPCPP-tubulin. To estimate the concentrations of GDP- and GMPCPP-tubulin for each nucleotide mixture, we assumed that only GMPCPP-tubulin contributes to minus-ended growth, consistent with our simulations (Figure 1 BC). This assumption allowed us to estimate the concentration of GMPCPP-tubulin in each nucleotide mixture by matching the observed growth rates to the control ‘all GMPCPP’ growth curve (Figure 2B). A potential problem with this approach is that the estimated GMPCPP-tubulin concentration in each mixture will be affected by error in the growth rate measurements. To minimize the impact of error, we performed a global fit to all measurements using a simple competitive inhibition model that enforced consistent nucleotide binding affinity (Figure 3A). The GMPCPP-tubulin concentrations that best recapitulate minus-end growth rates are consistent with tubulin binding 12.5-fold less tightly to GMPCPP (K_D_^GMPCPP^) than to GDP (K_D_^GDP^) (Figure 3B). This inferred difference of affinities was supported by direct measurements of nucleotide binding (Figure 3 – figure supplement 1), and agrees with prior reports (Correia et al., 1987; Hyman et al., 1992).

**Figure 3.**
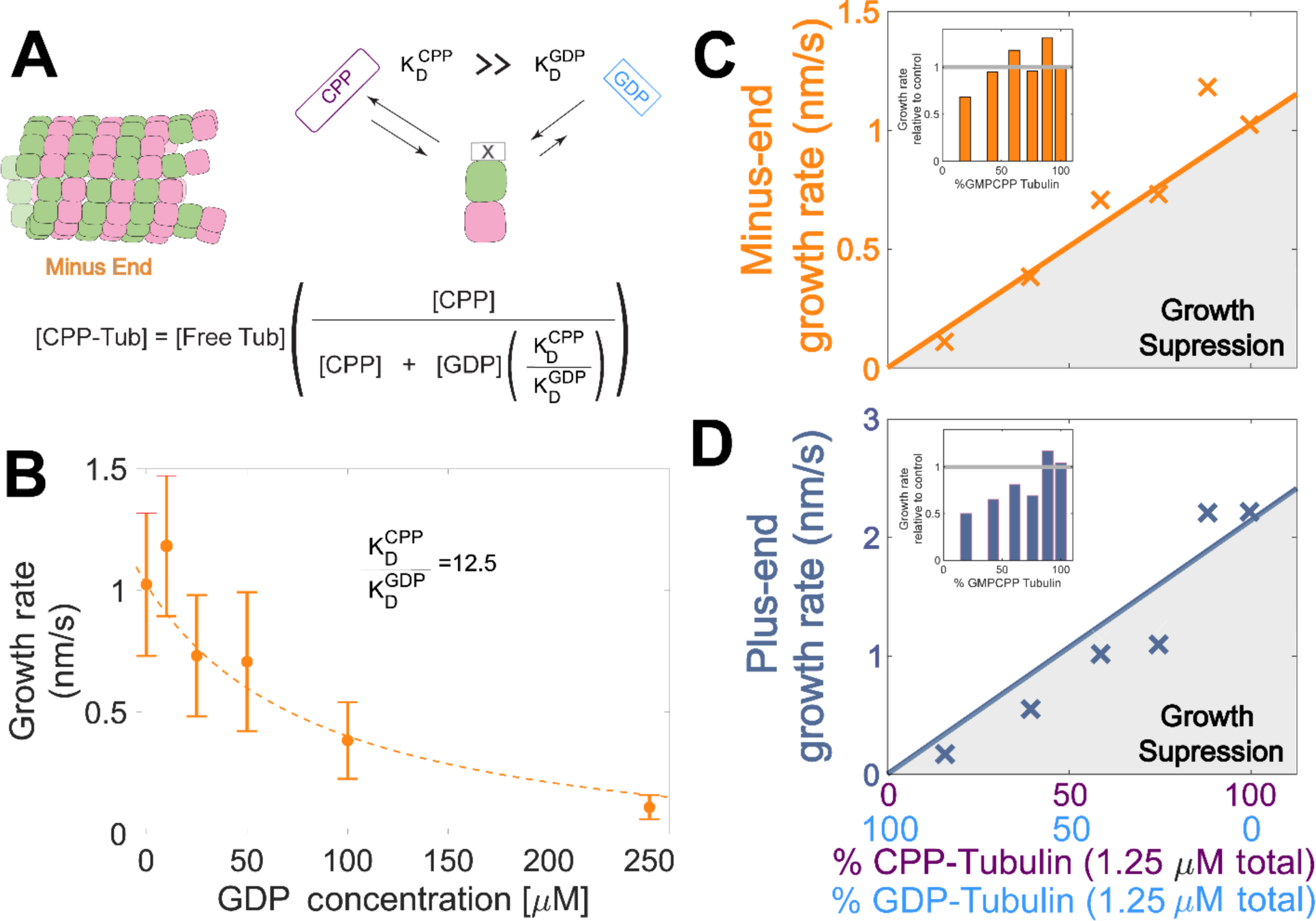
Microtubule plus-end growth is suppressed superstoichiometrically by GDP-tubulin. **(A)** Competitive nucleotide binding model. Mixed nucleotide assays result in either GDP- or GMPCPP-bound tubulin landing and creating a nucleotide interface at the minus-end. The concentration of GMPCPP-bound tubulin was determined using the concentrations of each nucleotide and their relative affinities (K_D_^CPP^/ K_D_^GDP^) through a competitive binding model (inset equation). **(B)** Minus-end growth rates. GMPCPP-tubulin was assumed to be the only tubulin that can contribute to minus-end growth in the mixed nucleotide assays. Minus-end growth rates over varying GDP concentrations were globally fit to a competitive inhibition model (Figure 3A), which resulted in a GMPCPP-tubulin concentration that was consistent with the ‘all-GMPCPP’ minus-growth curves (Figure 2B). The relative affinity of tubulin for GMPCPP compared to GDP (K_D_^CPP^/ K_D_^GDP^) was the only free parameter in the model. **(C-D)** Growth rates from Fig. 2C plotted as a function of the fraction of GDP-tubulin, estimated using the known nucleotide content and binding affinities. Growth rates are considered suppressed when falling below the solid lines exhibiting the ‘all-GMPCPP’ minus- and plus-end growth curves. Insets plot growth rates normalized to the ‘GMPCPP-only’ growth rates (gray solid line), showing a disproportionate decrease (∼1.5-fold for most concentrations) in plus-end growth. Differences in mean normalized growth rates at plus- and minus-ends were statistically significant with p < 0.001 for all nucleotide mixtures except 0% GDP (see Fig. 2).

To determine whether the observed decrease in growth rate was stoichiometric with the amount of GMPCPP-tubulin in the assay, we used the binding affinities and the known nucleotide concentrations in each mixture (Figure 3B) to extrapolate equivalent ‘GMPCPP-only’ growth rates (Figure 3 CD solid lines) from the control “all GMPCPP” curves (Fig 2A). Minus-end growth rates decreased stoichiometrically as the concentration of GDP-tubulin increased, matching or even slightly exceeding the “all GMPCPP” extrapolation. The only exception was at the highest concentration of GDP-tubulin (Figure 3C inset), where growth rates were slow and most challenging to quantify. In contrast, plus-end growth rates decreased super-stoichiometrically (were slower than expected based on the “all GMPCPP” extrapolation) for a given concentration of GDP-tubulin (Figure 3D). This super-stoichiometric effect at the plus-end was observed over a range of GDP-tubulin concentrations, and was most apparent when 25-55% of unpolymerized tubulin was bound to GDP (Figure 3D inset). The super-stoichiometric effect of GDP-tubulin on plus-end growth over a wide range of GDP-tubulin concentrations provides strong support for the interface-acting nucleotide mechanism.

### Why is plus-end growth hypersensitive to GDP-tubulin?

To establish a biochemical baseline for simulating mixed nucleotide states, we first fit the interface-acting nucleotide model to the “all GMPCPP” data (Fig. 2). The plus-end growth rates were recapitulated well using the same parameters obtained in a prior study ((Cleary et al., 2022); k_on_^plus^ of 0.74 µM^-1^ s^-1^, K_D_^longitudinal^ of 86 µM, K_D_^corner^ of 25 nM; Figure 4A). To extend the model to fit the minus-end growth rates, we retained the same interaction affinities as for the plus-end (consistent with the equal apparent critical concentration at both ends, Figure 2B) and optimized a minus-end-specific on-rate constant. This procedure yielded a k_on_^minus^ of 0.31 µM^-1^ s^-1^ (Figure 4A), roughly 2-fold slower than the plus-end, which is in line with the ∼2-fold lower concentration-dependence of growth rates observed at the minus-end (Figure 2A). We next performed mixed nucleotide (GMPCPP and GDP) simulations at a constant tubulin concentration of 1.25 µM. Simulated minus-end growth rates decreased linearly as the concentration of GDP-tubulin increased, recapitulating the experimental measurements (Figure 4B) and in agreement with our initial prediction using arbitrary parameters (Fig. 1). In contrast, simulated plus-end growth rates decreased super-stoichiometrically as the concentration of GDP-tubulin increased (Figure 4C). An outsized effect of GDP-tubulin at the plus-end is expected from the interface-acting mechanism, but the model overpredicted the magnitude of the effect.

**Figure 4.**
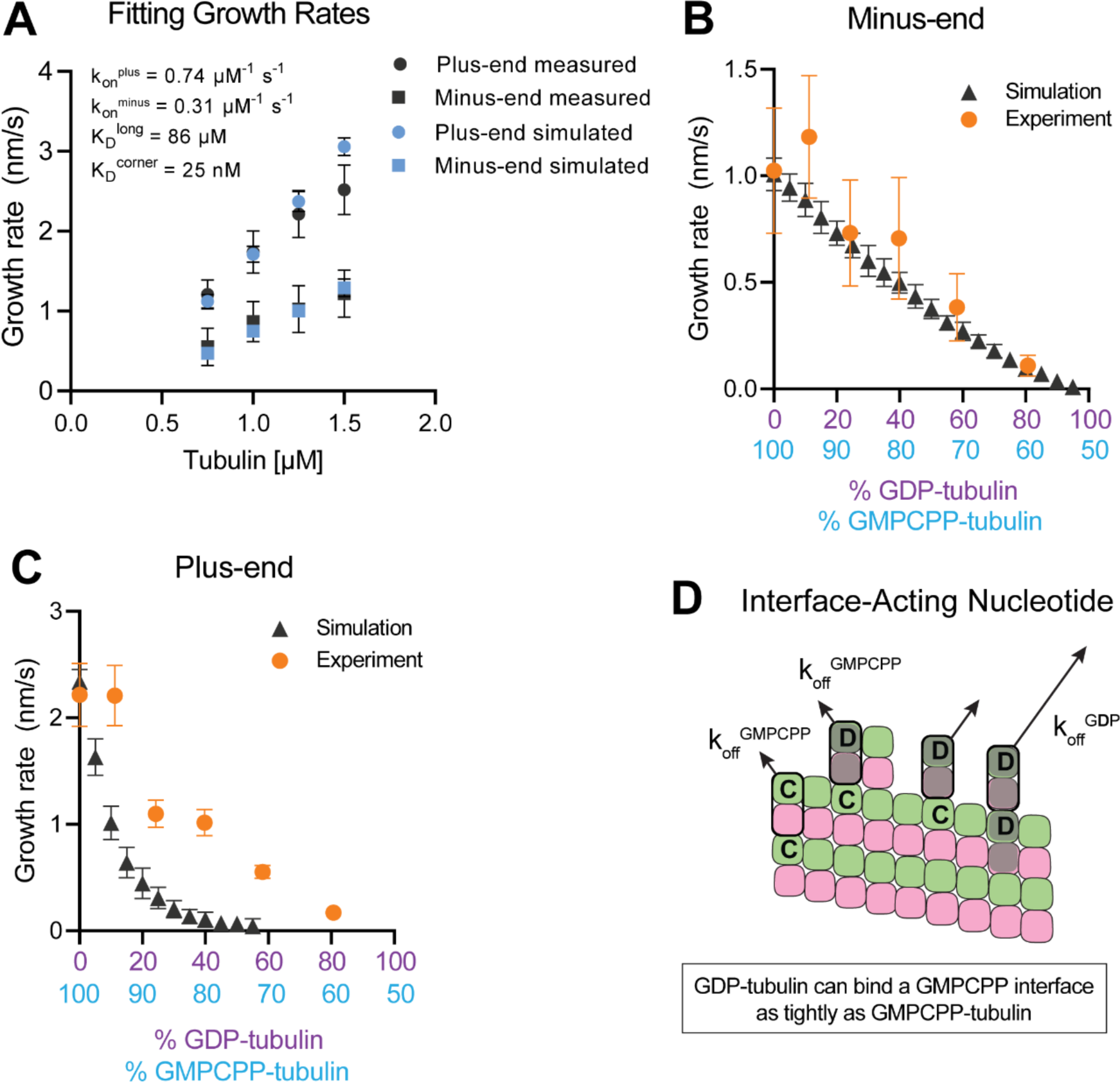
Simulating microtubule growth rates in the presence of GDP-tubulin. **(A)** Measured and simulated growth rates for plus- and minus-ends of GMPCPP microtubules. Inset shows the best-fit values for the plus-end and minus-end on-rate constants (k_on_^plus^ and k_on_^minus^, respectively), longitudinal interaction (K_D_^long^), and corner interaction (K_D_^corner^). Error bars show standard deviation (n = 50 per simulated concentration) and are obscured by symbols in some cases; experimental data are replotted from Figure 3). **(B and C)** Simulated and experimental growth rates at 1.25 µM tubulin in the presence of variable amounts of GDP-tubulin for microtubule minus-ends **(B)** and plus-ends **(C)**. **(D)** Cartoon showing how off-rates (k_off_) of GDP-tubulin at the plus-end are dependent upon the interfacial nucleotide; C = GMPCPP, D = GDP. Long-residing GDP-tubulin bound at corner- (one longitudinal and one lateral contact) or bucket-type (one longitudinal and two lateral contacts) binding sites explains the outsized effects of GDP-tubulin on plus-end elongation.

In the interface-acting nucleotide mechanism, the outsized effect of GDP at the plus-end occurs because GDP-tubulin can bind tightly to (and reside longer at) the plus-end if the interfacial nucleotide is GMPCPP (Figure 4D). This ‘extended stay’ of GDP-tubulin on the plus-end poisons the protofilament end against further growth and is the origin of the super-stoichiometric effect of GDP-tubulin at the plus-end (Figure 4D). We reasoned that our model might be overpredicting the magnitude of the GDP-poisoning effect (Figure 4C) because it was neglecting some other mechanism that normally limits the lifetime of GDP-tubulin at the plus-end.

### Nucleotide exchange at the plus-end can alleviate protofilament ‘poisoning’ by GDP-tubulin

Recent work (Luo et al., 2023; Piedra et al., 2016) has reinforced early results (Chen & Hill, 1983; Chen & Hill, 1985; Mitchison, 1993) that pointed to the potential role of nucleotide exchange in microtubule dynamics at the plus-end. We implemented a finite rate of nucleotide exchange in the model (see Methods) to determine whether exchange might allow the simulations to better recapitulate the magnitude by which GDP-tubulin super-stoichiometrically decreased plus-end growth (Figure 5A). We performed interface-acting plus-end simulations using a range of nucleotide exchange rates (Figure 5 – figure supplement 1). Faster rates of nucleotide exchange yielded smaller decreases in plus-end growth rates for a given concentration of GDP-tubulin (Figure 5B). The rate of nucleotide exchange that best recapitulated the observed effects was 0.3 – 0.5 s^-1^, which compares favorably to other estimates (Amayed et al., 2000; Melki et al., 1989; Yarbrough & Fishback, 1985) (Figure 5 – figure supplement 1). In summary, using simulations and measurements of plus- and minus-end growth, we showed that microtubule plus- and minus-ends exhibit different sensitivities to GDP-tubulin, lending strong support for the interface-acting mechanism of nucleotide action.

**Figure 5.**
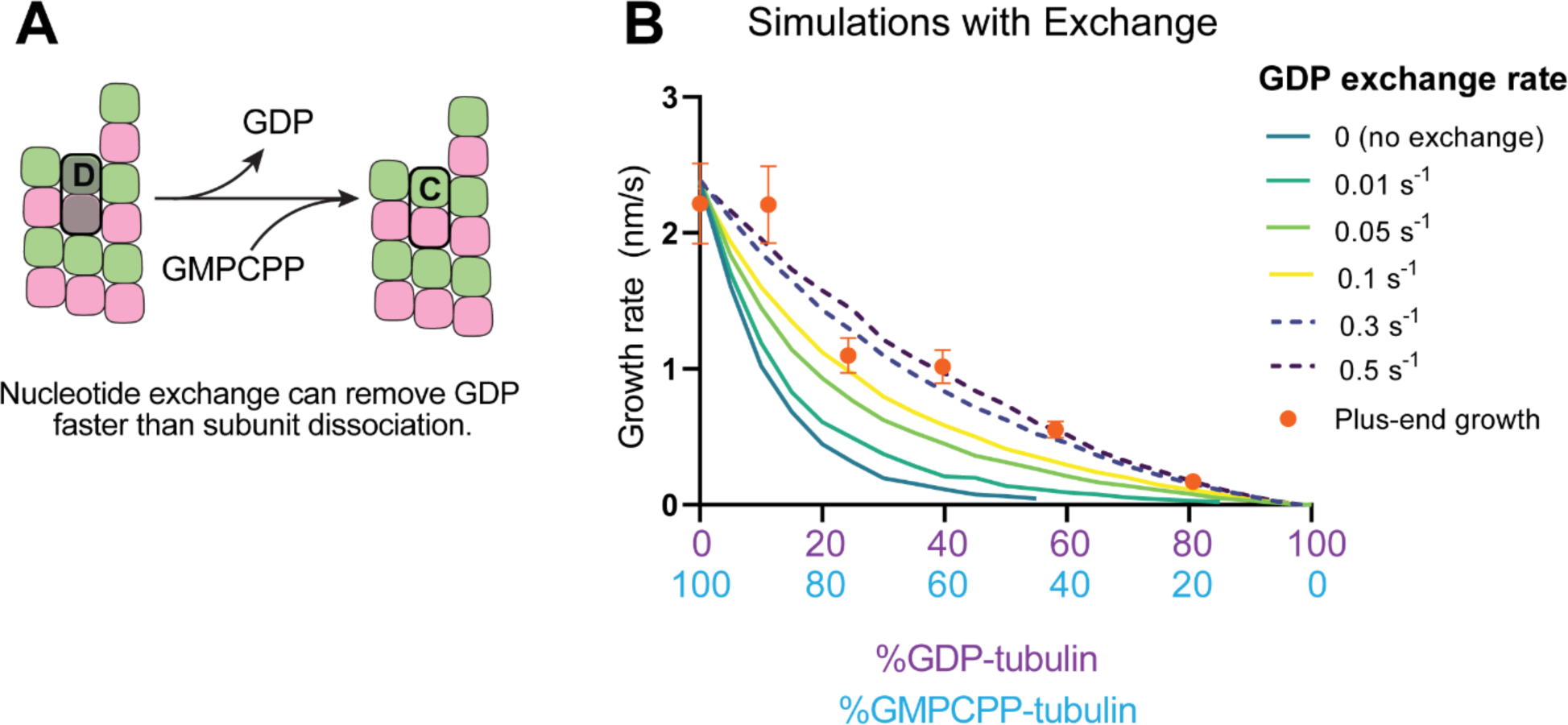
Effects of nucleotide exchange on simulated microtubule plus-end growth rates. **(A)** Nucleotide exchange on terminal subunits can mitigate protofilament poisoning at microtubule plus-ends by reducing the lifetime of GDP on the microtubule end. **(B)** Simulated growth rates of microtubule plus-ends as a function of the nucleotide exchange rate (see Methods), showing that faster rates of exchange modulate the effect of protofilament poisoning. Orange circles show the measured plus-end growth rates (Figure 3).

## Discussion

A connection between tubulin nucleotide state and microtubule stability has long been appreciated, but the molecular mechanism underlying the connection has been surprisingly difficult to determine. At one extreme, a self-acting mechanism inspired by conformational differences between unpolymerized and polymerized tubulin posits that GTP dictates microtubule stability by promoting a more microtubule-compatible conformation for the tubulin *to which it is bound*. At the other extreme, an interface-acting mechanism inspired by direct participation of nucleotide in tubulin:tubulin polymerization contacts posits that the nucleotide influences the behavior of the *next* tubulin along the protofilament. Ruling out either the self- or interface-acting mechanism has been challenging because it has not been possible to manipulate the nucleotide on the plus-end separately from the nucleotide on unpolymerized tubulin. Consequently, tests to date have relied on indirect data.

In the present study, we took advantage of microtubule polarity to address the debate about the mechanism of nucleotide action in a new way. Our approach rests on an asymmetry in the way that nucleotide participates in plus- and minus-end interactions. At the minus-end there is no difference between self- and interface-acting nucleotide mechanisms because the nucleotide on the terminal tubulin is also the interfacial nucleotide that participates in contacts along the protofilament. At the plus-end, however, two different nucleotide binding sites are involved: the one exposed on the terminal tubulin, and one at the interface with (underneath) the terminal tubulin. Our computational simulations of microtubule elongation revealed that the two mechanisms make different predictions about the sensitivity of plus- and minus-end elongation to GDP-tubulin. We used mixed nucleotide (GMPCPP and GDP) experiments to measure the effects of GDP-tubulin on elongation of plus- and minus-ends in a way that controls nucleotide state(s) while also avoiding complications associated with microtubule catastrophe.

We observed that the decrease in elongation rate was proportionally greater for the plus-end than for the minus-end over a wide range of GDP-tubulin fractions. This outsized effect at the plus-end is consistent with an earlier study that observed a loss of plus-end elongation when using mixtures of GTP and GDP (Tanaka-Takiguchi et al., 1998). The outsized effect at the plus-end conforms to predictions of the interface-acting mechanism and is incompatible with the self-acting mechanism. To recapitulate the magnitude of GDP-tubulin-induced suppression of growth rates, the simulations required a finite rate of nucleotide exchange on plus-end protofilaments. Our experiments and simulations provide strong new data that support the interface-acting mechanism of nucleotide action.

The outsized effect of GDP on plus-ends provides new insights into the fundamental mechanisms of microtubule dynamics and adds to a growing body of evidence that suggests GDP-terminated protofilaments influence microtubule growth (Carlier & Pantaloni, 1978; Hamel et al., 1986; Valiron et al., 2010), fluctuations (Cleary et al., 2022), catastrophe (Caplow & Shanks, 1996; Piedra et al., 2016), and regulation (Lawrence et al., 2022; Luo et al., 2023). Our results point to a more nuanced view of the GTP cap model, which posits that growing microtubule ends are protected against depolymerization by a ‘cap’ of GTP-tubulin ((Mitchison & Kirschner, 1984), reviewed in (Gudimchuk & McIntosh, 2021)). Early views of the GTP cap did not anticipate the influence of GDP-tubulin on growing plus-ends, but there is now increasing evidence (Carlier & Pantaloni, 1978; Farmer & Zanic, 2023; Hamel et al., 1986; Margolin et al., 2012; Maurer et al., 2012; Roth et al., 2018; Valiron et al., 2010) that the cap is not ‘all or nothing’, and that GDP-tubulin can modulate microtubule growth without always initiating a catastrophe. Indeed, the tendency for plus-ended growth to ‘stutter’ (Mahserejian et al., 2022) and fluctuate (Cleary et al., 2022) might be explained by exposed GDP-tubulin; exposed GDP-tubulin may also contribute to the higher frequency of catastrophe at the plus-end (Strothman et al., 2019; Walker et al., 1988). Our work supports an emerging view of the growing microtubule end as a ‘mosaic’ of nucleotide states rather than a uniform assembly of GTP-tubulin (Brouhard & Sept, 2012; Brouhard & Rice, 2018; Cross, 2019; Duellberg et al., 2016; Farmer et al., 2021; Farmer & Zanic, 2023; Gudimchuk & McIntosh, 2021; Howard & Hyman, 2009; Margolin et al., 2012; Maurer et al., 2012; Roostalu et al., 2020; Roth et al., 2018). By allowing for the possibility of multiple nucleotide states on the microtubule end, our work also resonates with recent studies of the microtubule regulatory factor CLASP (Lawrence et al., 2022; Luo et al., 2023), which regulates microtubule plus-ends differently depending on the nucleotide state of the terminal subunit at the protofilament plus-end.

Our modeling purposefully implemented the simplest forms of self- and interface-acting nucleotide mechanisms. We did not attempt to explicitly model how conformations of αβ-tubulin might influence the strength of tubulin:tubulin interactions: there is no consensus about how to do so, and modeling different conformations introduces substantially more adjustable parameters into the model, which complicates fitting and interpretation (Coombes et al., 2013; Molodtsov et al., 2005; Stewman et al., 2020; VanBuren et al., 2005; Zakharov et al., 2015). In support of a simpler model, our use of GMPCPP and GDP mixtures simplified the biochemical picture by ensuring that, except for the very end, the microtubule lattice will be predominantly in a single nucleotide state (GMPCPP). This choice diminishes the importance of explicitly modeling different conformations. The model might also implicitly capture a subset of the conformation-dependent effects on tubulin:tubulin interfaces because the longitudinal and corner affinities are refined independently: the strength of longitudinal interactions could therefore be different for corner than for pure longitudinal sites, potentially reflecting the cost of tubulin ‘straightening’ during polymerization. In the interest of minimizing the number of adjustable parameters in the model, we also did not consider ‘hybrid’ models incorporating elements from both self- and interface-acting mechanisms. While we acknowledge the possibility that self-acting mechanisms may contribute to modulation of plus-end stability, the large differences we predicted and observed between plus- and minus-ends indicate that interface-acting nucleotide effects are sufficient to explain the observations. This interface-centric view of nucleotide action is also consistent with cryo-EM studies, which show that the largest nucleotide-dependent conformational changes in the microtubule occur in the α-tubulin subunit above and directly contacting the β-tubulin exchangeable nucleotide (Alushin et al., 2014; Manka & Moores, 2018; Zhang et al., 2015).

In summary, the findings reported here provide the most direct evidence to date in support of an interface-acting mechanism for nucleotide in microtubule stabilization. Depolymerizing microtubules can also perform mechanical work, and it is interesting to consider parallels with other work-performing, oligomeric nucleotide hydrolases. The curling protofilaments that occur during microtubule depolymerization are effectively linear oligomers held together (and to the microtubule end) by nucleotide-dependent interactions at the tubulin:tubulin interfaces. AAA-family proteins, which are oligomeric ATPases that use nucleotide-dependent reorganization of quaternary structure to unfold proteins, package viral DNA, and remodel the structure of nucleic acids (Banerjee et al., 2016; Brunger & DeLaBarre, 2003; Davies et al., 2005; Erzberger & Berger, 2006), also appear to use an interface-acting mechanism for their bound adenosine nucleotide. Indeed, just as the GTP-binding site on β-tubulin forms part of the longitudinal interface between tubulin subunits, the ATP binding site in AAA proteins resides at a protomer:protomer interface and dictates the geometry of oligomerization contacts (Erzberger & Berger, 2006). Furthermore, for both AAA proteins and microtubules, residues important for nucleotide hydrolysis on one subunit are contributed by the next subunit in the oligomer or polymer (Banerjee et al., 2016; Brunger & DeLaBarre, 2003; Davies et al., 2005). We speculate that these similarities involving interfacial nucleotides in otherwise unrelated proteins may indicate a shared, convergently evolved mechanism for achieving force production in oligomers.

## Acknowledgements

This study was supported by NIH R01-GM135565 to LMR, and by NIH R35-GM139568 to WOH. JMC received support from NIH-T32 GM108563, and LAM received support from a Postdoctoral Research Fellowship in Biology from the NSF (Award 2209298).

## Methods

### Protein purification and labeling

PC-grade bovine brain tubulin was purified as previously described (Cleary et al., 2022; Uppalapati et al., 2009), double cycled, quantified by absorbance at 280 nm (ε_tubulin_ of 115,000 M^-1^cm^-1^), diluted to 100 µM in BRB80 (80 mM K-Pipes, 2 mM EGTA, 2 mM MgCl_2_, pH 6.9), aliquoted, flash frozen in liquid nitrogen, and stored at -80°C. Prior to experiments, tubulin aliquots were thawed on ice, diluted to 20 µM in BRB80, and concentrations reconfirmed by A_280_.

Tubulin was biotinylated as previously described (Cleary et al., 2022). Briefly, microtubules were polymerized by combining 40 µM tubulin, 1 mM GTP, 1 mM MgCl_2_ and 5% DMSO in BRB80, incubating at 37°C for 30 minutes. An equimolar amount of EZ-Link NHS-Biotin in DMSO (ThermoFisher 20217) was added and allowed to react for 30 minutes at 37°C. Microtubules were then pelleted, the pellet resuspended in cold BRB80 and incubated on ice for 30 minutes to depolymerize the microtubules, the solution centrifuged at 30 psi for 10 minutes in a Beckman Airfuge using a pre-chilled rotor, and supernatant collected. This biotinylated tubulin was then cycled, the tubulin concentration checked by A_280_, and the degree of biotinylation quantified using the Biocytin Biotin Quantification Kit (Thermo Scientific #44610). Final stocks of biotinylated tubulin were mixed with unlabeled tubulin to 40 µM total tubulin to obtain a 33% biotin-labeled fraction, aliquoted, frozen in liquid nitrogen, and stored at -80°C.

Biotinylated microtubule seeds were polymerized by combining 20 µM biotinylated tubulin (33% biotin-labeled), 1 mM GMPCPP (Jena Biosciences) and 4 mM MgCl_2_, and incubating at 37 °C for 1 hour. The seeds were then elongated by diluting the total tubulin concentration to 2 µM in BRB80 with 0.5 mM GMPCPP and 2 mM MgCl_2_ and incubating for 5 hours at 37 °C. The seeds were pelleted, resuspended in BRB80 with 20% glycerol, flash frozen in liquid nitrogen, and stored at -80°C. On the day of experiments the aliquot was rapidly thawed at 37 °C, the seeds pelleted to remove glycerol, and resuspended in a solution containing 0.5 mM Mg-GMPCPP.

### Microtubule dynamics assays

Coverslips (18 × 18 mm Corning) were cleaned in 7X Cleaning Detergent (MP Biomedicals™ 097667093) diluted to 1X in ddH_2_O. The solution was heated at 45°C until clear, the coverslips were then immersed for 2 hours, removed and rinsed with ddH_2_O, and plasma cleaned (Harrick Plasma) for 12 minutes. Following cleaning, coverslips were silanized by incubating in a vacuum-sealed desiccator with 1H,1H,2H,2H-perfluorodecyltrichlorosilane (Alfa Aesar L165804-03) overnight. Before use, the degree of silanization was checked using a droplet test to confirm hydrophobicity.

To construct flow cells, a second ethanol-washed and ddH_2_O-rinsed coverslip (60 x 24 mm Corning) was scored, split to a width smaller than 18 mm, and attached to the silanized coverslip with two strips of double-sided tape spaced roughly 10 mm apart. For the experiment, 600 nM neutravidin (ThermoFisher) was flowed into the chamber, followed by 5% F127 (Sigma P2443-250G), 2 mg/mL casein (Sigma C-7078), and biotinylated microtubule seeds at a concentration that resulted in approximately 10 seeds per 90 x 90 µm^2^ field of view. Biotinylated BSA (1 mg/mL) was then added to the flow chamber to block any free neutravidin on the cover slip.

Due to the slow growth conditions in these experiments, it was necessary to pre-establish the plus- and minus-ends of the seeds in every field of view. Polarity was determined by injecting into the flow cell a solution containing 12.5 µM tubulin, 1 mM Mg-GTP and an oxygen scavenging system consisting of 80 μg/mL Catalase [Sigma C1345-1G], 100 mM DTT, 200 mM D-Glucose [EMD Millipore Corp DX0145-1], and 200 μg/mL Glucose Oxidase [EMD Millipore Corp 345386–10gm] in BRB80). The flow cell was allowed to warm for 5 minutes in contact with the objective of the Nikon TE-2000 TIRF with an objective heater set to 30°C. Microtubules were visualized using IRM with a blue (440 nm) LED at 0.5 % power (pE-300white, CoolLED, UK). Microtubule growth in GTP was monitored for 5 minutes, and the faster growing end of each microtubule in the field was defined as the plus-end. The tubulin solution was then replaced with tubulin-free cold BRB80, the flow cell was incubated for 5 minutes to depolymerize the microtubules with depolymerization confirmed by visualization, and finally any residual tubulin was removed by flowing through five flow cell volumes of cold BRB80.

While monitoring the same field of view, a solution was introduced containing tubulin, 1 mM nucleotide (either Mg-GMPCPP or a mixture of Mg-GDP/GMPCPP, with concentrations quantified by absorbance at 252 nm, using ε = 13,700 M^-1^ cm^-1^), and an oxygen scavenging system. Once the final polymerization mixture was introduced, the chamber was sealed with nail polish and allowed to equilibrate to 30°C while in contact with the objective. Images were subsequently taken at 1 frame per second for up to 2.5 hours.

### Image analysis and processing

Each video was flat-fielded to correct for uneven illumination, as follows using ImageJ (Schindelin et al., 2015). First, an out-of-focus movie was acquired and a median image generated. The median image was then converted to 32-bits, and normalized to 1 by dividing every pixel value by the mean pixel intensity in the image. Finally, every experimental video was flat-fielded by dividing the intensity values in every frame by this normalized median image. Stage drift was corrected as previously described (Cleary et al., 2022): fiduciary markers were tracked using FIESTA (Ruhnow et al., 2011) and used as input for an in-house drift correction program written in Matlab. To quantify microtubule growth, kymographs were generated from pixel-corrected movies using the line-scan tool in ImageJ. Plus- and minus-end growth rates were determined by fitting a line to smooth and continuous growth events and calculating the slope.

### Global fit of minus-end growth

The relative binding affinities of tubulin for GDP and GMPCPP were estimated by fitting the minus-end growth rates at varying nucleotide ratios to a model in which only GMPCPP-tubulin contributes to minus-end growth, as follows. From Fig. 2B, the minus-end growth rate (GR_minus_ in nm/s) as a function of [tubulin_GMPCPP_] (in µM) was:

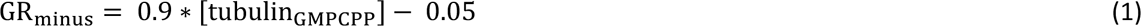

In a mixture of GMPCPP and GDP, the concentration of GMPCPP-tubulin can be determined by a competitive binding model (analogous to competitive inhibition of an enzyme (Cheng & Prusoff, 1973)) in which the two nucleotides compete for binding to tubulin:

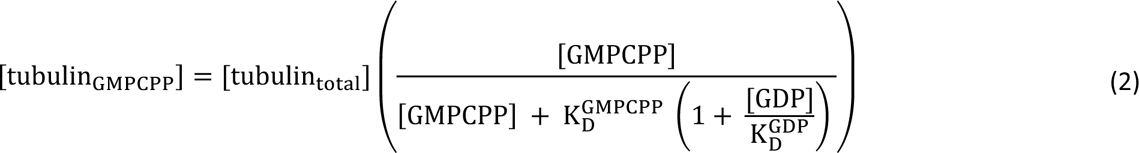

Because [GMPCPP] was relatively high in all cases, we made the assumption that 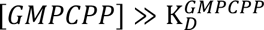, which simplifies Eq. (2) to:

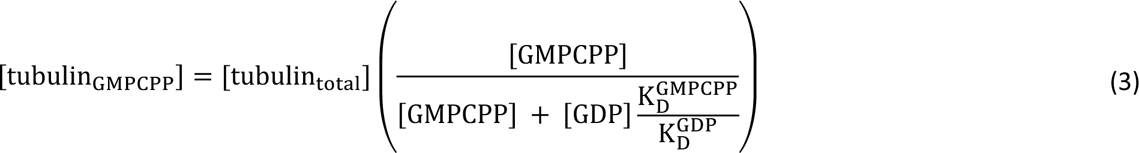

Plugging (3) into (1) gives:

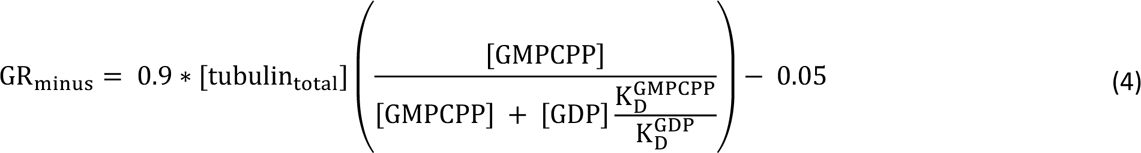

Finally, because the total nucleotide concentration was kept constant at 1000 µM, we could replace [GMPCPP] by 1000 – [GDP], yielding:

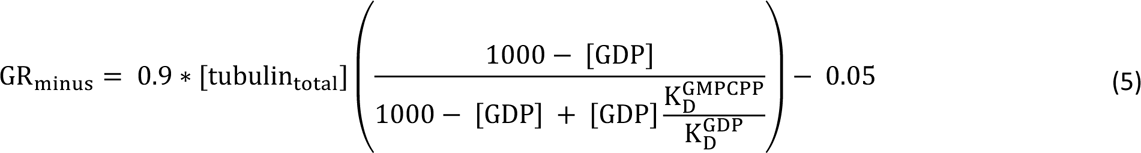

The minus-end growth rates as a function of [GDP] in Fig. 2B (where [tubulin_*+*,-_] was 1.25 µM) were fit to Eq. (5). Here, the only free parameter is the relative affinity of tubulin for GMPCPP and GDP (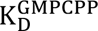/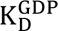). The fit was weighted by the inverse of the standard error of the mean (SEM).

### Nucleotide binding affinity assays

The affinity of tubulin for GMPCPP and GDP was determined using a competition assay that relies on the quenching of tryptophan fluorescence by 6-Thio GTP (Amayed et al., 2000; Fishback & Yarbrough, 1984; Piedra et al., 2016). Aliquots of tubulin (∼ 80 μM) and Bovine Serum Albumin (BSA – 50 mg/mL) were rapidly thawed, filtered through a 0.1 µm spin filter (Millipore-Sigma, UFC30VV25) at 11,000 rpm and 4°C to remove aggregates, and concentrations quantified by absorbance at A_280_ with ε_tubulin_=115,000 M^-1^ cm^-1^ and ε_BSA_=43,824 M^-1^ cm^-1^. The affinity of 6-Thio GTP for tubulin was measured by preparing 220 μL samples of either 0.2 μM tubulin or 0.56 μM BSA with varying concentrations of 6-Thio GTP. The BSA concentration was chosen to match the tryptophan fluorescence of tubulin, which allowed for the correction of the inner filter effect due to absorbance of 6-Thio GTP at the tryptophan emission peak. A buffer-only well was included in every plate as a zero fluorescence control, and the value of the blank was subtracted from each BSA and tubulin measurement. Tryptophan fluorescence readings (297 nm excitation and 332 nm emission) were performed in 96-well, flat bottom, UV-star plates (Greiner bio-one, 655809) on a Molecular Devices FlexStation 3 Multimode Microplate Reader. Each recorded fluorescence value was an average of 250 signal determinations. The fluorescence readings were corrected for the inner filter effect by dividing the tubulin fluorescence signal by the BSA fluorescence signal at each nucleotide concentration (Fishback and Yarbrough, 1984). The standard error of the mean (SEM) was calculated using propagated errors of the relative SEM for each variable:

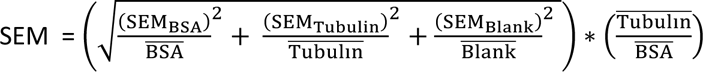

The affinity of tubulin for 6-Thio GTP (K_D_^6-Thio GTP^) was determined by adding increasing concentrations of 6-Thio GTP, measuring the fall in fluorescence due to fluorescence quenching by the nucleotide, and fitting the data to a binding isotherm weighted by the inverse of the SEM:

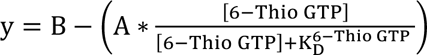

where A corresponds to the amplitude of the fall in fluorescence and (B - A) is the remaining fluorescence under full quenching conditions. Competition assays were then performed by adding increasing concentrations of GMPCPP or GDP to a solution containing 3 μM 6-Thio GTP. Competition between the unlabeled nucleotides and the 6-Thio GTP caused unquenching of fluorescence, allowing for determination of the affinity of tubulin for GMPCPP (K_D_^GMPCPP^) and GDP (K_D_^GDP^). Data for each nucleotide were fit to a competition model:

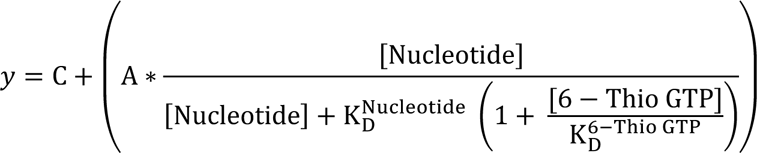

Here, A is the amplitude fluorescence quenching, which was constrained by the measured value at 3 μM 6-Thio GTP, and C is a free parameter corresponding to the quenched fluorescence value at zero unlabeled nucleotide. For this fit, the means were weighted by the inverse of the SEM.

### Simulating microtubule growth of plus- and minus-ends

Simulations of plus- and minus-end elongation were performed using extended versions of previously-described code and analysis algorithms (https://git.biohpc.swmed.edu/s422146/simulate-mt-44 (Rice, 2022)) (Cleary et al., 2022; Kim & Rice, 2019; Mickolajczyk et al., 2019; Piedra et al., 2016). The main features of the model are outlined in Supplemental Figure 1 and described in detail in our previous publication (Cleary et al., 2022). Briefly, the code performs kinetic Monte Carlo simulations of microtubule elongation at the level of individual association and dissociation events, creating a ‘biochemical movie’ of polymerization with one reaction (association, dissociation, or nucleotide exchange) per frame. First-order subunit association rate constants are calculated by multiplying the bimolecular on-rate constant (k_on_) by the tubulin concentration (on-rate = k_on_*[tubulin]). Subunit dissociation rates (k_off_ = k_on_*K_D_) are dependent on the interaction affinity (K_D_) at a specific site, which is determined by the number and type of tubulin-tubulin interactions at the respective site.

The simulation code from the present work is available as a GitLab repository (https://git.biohpc.swmed.edu/ricelab/simulate-mt-52, (Rice, 2023)). To compare the effects of interface-acting and self-acting mechanisms, plus-end simulation code was modified to implement self-acting (cis) nucleotide instead of interface-acting nucleotide (Figure 1 Supplement 1B). To simulate the minus-end, simulation rules were updated to reflect the lack of an exposed nucleotide on the minus-end, and the orientation of interactions across the seam (Figure 1 Supplement 1C). Simulating GDP- and GMPCPP-tubulin mixtures required two new parameters: (1) the concentration of GDP-tubulin and (2) a factor to weaken GDP-mediated contacts, represented as a multiplicative factor on the GMPCPP interaction affinity. We assumed that there were no inherent differences between the association rates of GMPCPP- and GDP-tubulin, and used the same on-rate constant (k_on_^plus^ or k_on_^minus^, respectively) for GDP- and GMPCPP-tubulin.

We generalized our prior implementation of nucleotide exchange (Piedra et al., 2016) to allow all terminal nucleotides (whether GMPCPP or GDP) to exchange. The probability of replacement by GDP or GMPCPP was set to be proportional to the fractional concentration of either nucleotide. Our implementation assumes that the rate-limiting step in nucleotide exchange is dissociation of the (previously) bound nucleotide. To reflect the 12.5-fold difference between the affinity of tubulin for GDP and GMPCPP, the rate of GMPCPP exchange (the off-rate) was set to be 12.5-fold faster than the rate of GDP exchange (as measured in Fig. 3).

### Constraining biochemical parameters for simulations

Experimental plus-end growth rates were recapitulated using biochemical parameters obtained in a prior study (k_on_^plus^ = 0.74 µM^1^ s^-1^, K_D_^longitudinal^ = 86 µM, K_D_^corner^ = 25 nM) (Cleary et al., 2022). Simulations of the minus-end used the same K_D_^longitudinal^ and K_D_^corner^ as for the plus-end. Iterative fitting in MATLAB was used to optimize an on-rate constant (k_on_^minus^) that could best recapitulate experimentally observed minus-end growth rates. Each fitting attempt used 50 independent simulations, 300 seconds in length, of minus-end growth at the same concentrations used for measurements of GMPCPP microtubules.

For all other simulations, 50 independent simulations of 600 seconds were run for each condition tested. Mixed nucleotide simulations in Figure 1 were performed at 1 µM total [αβ-tubulin], with varying percentages of GDP-tubulin (up to 20%, in 5% increments). Mixed nucleotide simulations in Figures 4 and 5 were performed at 1.25 µM total [αβ-tubulin] to mimic experimental conditions, with varied percentages of GDP-tubulin (up to 50%, in 5% increments). Simulations with GDP-tubulin used a ‘GDP weakening factor’ comparable to one used previously (Cleary et al., 2022), such that the GDP longitudinal interface was 3000- (in Fig. 1, where we used arbitrary affinities) or 3,500-fold weaker (in Figs. 4 and 5, where we fit affinities to recapitulate growth rates) than the GMPCPP-longitudinal interface; this magnitude weakening is consistent with the large difference between depolymerization rates of GMPCPP and GDP microtubules. Additional simulations in Figure 1 Supplements 2 and 3 used a GDP weakening factor such that the GDP longitudinal interface was 300-fold or 30,000-fold weaker than the GTP longitudinal interface.

Simulation parameters and calculated rates for Figures 1 – 5 are summarized in Tables 1 and 2. Parameters and calculated rates for Supplemental Figures are summarized in Tables 3 and 4.

**Table 2.** Calculated on-rates and off-rates for simulations presented in figures 1 – 5. On-rate is calculated using the biochemical k_on_ and the concentration of tubulin [µM]. Off-rate is calculated using the biochemical k_on_ and the dissociation constant K_D_.

| | Simulated end | On-rate<br>($s^{-1}$) | $k_{off}^{long}$<br>($s^{-1}$) | $k_{off}^{corner}$<br>($s^{-1}$) | $k_{off}^{long} GDP$ ($s^{-1}$) | $k_{off}^{corner} GDP$ ($s^{-1}$) | Tubulin<br>($\mu M$) |
| --- | --- | --- | --- | --- | --- | --- | --- |
| **Fig. 1** | plus | 1 | 100 | 0.1 | $3 \times 10^5$ | 300 | 1 |
| | minus | 1 | 100 | 0.1 | $3 \times 10^5$ | 300 | 1 |
| **Fig. 4A** | plus | 0.9 | 64 | 0.019 | $2.2 \times 10^5$ | 65 | 1.25 |
| | minus | 0.4 | 27 | 0.0078 | $9.3 \times 10^4$ | 27.3 | 1.25 |
| **Fig. 4B** | plus | 0.9 | 64 | 0.019 | $2.2 \times 10^5$ | 65 | 1.25 |
| **Fig. 4C** | minus | 0.4 | 27 | 0.0078 | $9.3 \times 10^4$ | 27.3 | 1.25 |
| **Fig. 5B** | plus | 0.9 | 64 | 0.019 | $2.2 \times 10^5$ | 65 | 1.25 |

**Table 3.** Simulation parameters for Figure 1 supplements. Shading has been added to highlight which simulation parameters were changed, with respect to the reference parameters used in Figure 1. The GDP fold weaker values are the fold change between the GTP- and GDP-type interaction.

| Fig. 1<br>Supp. 3 | change | $k_{on}$<br>($\mu M^{-1} s^{-1}$) | $K_D^{long}$<br>GTP<br>($\mu M$) | $K_D^{corner}$<br>GTP<br>($\mu M$) | $K_D^{long}$<br>GDP<br>($\mu M$) | $K_D^{corner}$<br>GDP<br>($\mu M$) | GDP<sup>long</sup><br>fold<br>weaker | GDP<sup>corner</sup><br>fold<br>weaker | Tubulin<br>($\mu M$) |
| --- | --- | --- | --- | --- | --- | --- | --- | --- | --- |
| A | GDP weakening | 1.0 | 100 | 0.1 | $3 \times 10^6$ | 3000 | 30,000 | 30,000 | 1 |
| A | GDP weakening | 1.0 | 100 | 0.1 | $3 \times 10^5$ | 300 | 3,000 | 3,000 | 1 |
| A | GDP weakening | 1.0 | 100 | 0.1 | $3 \times 10^4$ | 30 | 300 | 300 | 1 |
| B | $K_D^{long}$ | 1.0 | 1000 | 0.1 | $3 \times 10^6$ | 300 | 3,000 | 3,000 | 1 |
| B | $K_D^{long}$ | 1.0 | 100 | 0.1 | $3 \times 10^5$ | 300 | 3,000 | 3,000 | 1 |
| B | $K_D^{long}$ | 1.0 | 10 | 0.1 | $3 \times 10^4$ | 300 | 3,000 | 3,000 | 1 |
| C | $K_D^{corner}$ | 1.0 | 100 | 0.5 | $3 \times 10^5$ | 1500 | 3,000 | 3,000 | 1 |
| C | $K_D^{corner}$ | 1.0 | 100 | 0.1 | $3 \times 10^5$ | 300 | 3,000 | 3,000 | 1 |
| C | $K_D^{corner}$ | 1.0 | 100 | 0.02 | $3 \times 10^5$ | 60 | 3,000 | 3,000 | 1 |

**Table 4.** Calculated on-rates and off-rates for simulations in Figure 1 supplements. On-rate is calculated using the biochemical k_on_ and the concentration of tubulin [µM]. Off-rate is calculated using the biochemical k_on_ and the dissociation constant K_D_. All simulations in Figure 1 Supplement 2 and 3 use the same biochemical k_on_ for plus-end and minus-end simulations, as was done in Figure 1. Shading highlights which off-rates changed, with respect to the original values in Figure 1.

| Fig. 1<br>Supp. 3 | On-rate ($s^{-1}$) | $k_{off}^{long}$ GTP<br>($s^{-1}$) | $k_{off}^{corner}$ GTP<br>($s^{-1}$) | $k_{off}^{long}$ GDP<br>($s^{-1}$) | $k_{off}^{corner}$ GDP<br>($s^{-1}$) | Tubulin<br>($\mu M$) |
| --- | --- | --- | --- | --- | --- | --- |
| A | 1 | 100 | 0.1 | $3 \times 10^6$ | 3000 | 1 |
| A | 1 | 100 | 0.1 | $3 \times 10^5$ | 300 | 1 |
| A | 1 | 100 | 0.1 | $3 \times 10^4$ | 30 | 1 |
| B | 1 | 1000 | 0.1 | $3 \times 10^6$ | 300 | 1 |
| B | 1 | 100 | 0.1 | $3 \times 10^5$ | 300 | 1 |
| B | 1 | 10 | 0.1 | $3 \times 10^4$ | 300 | 1 |
| C | 1 | 100 | 0.5 | $3 \times 10^5$ | 1500 | 1 |
| C | 1 | 100 | 0.1 | $3 \times 10^5$ | 300 | 1 |
| C | 1 | 100 | 0.02 | $3 \times 10^5$ | 60 | 1 |

---

**Figure 1 - figure supplement 1.**
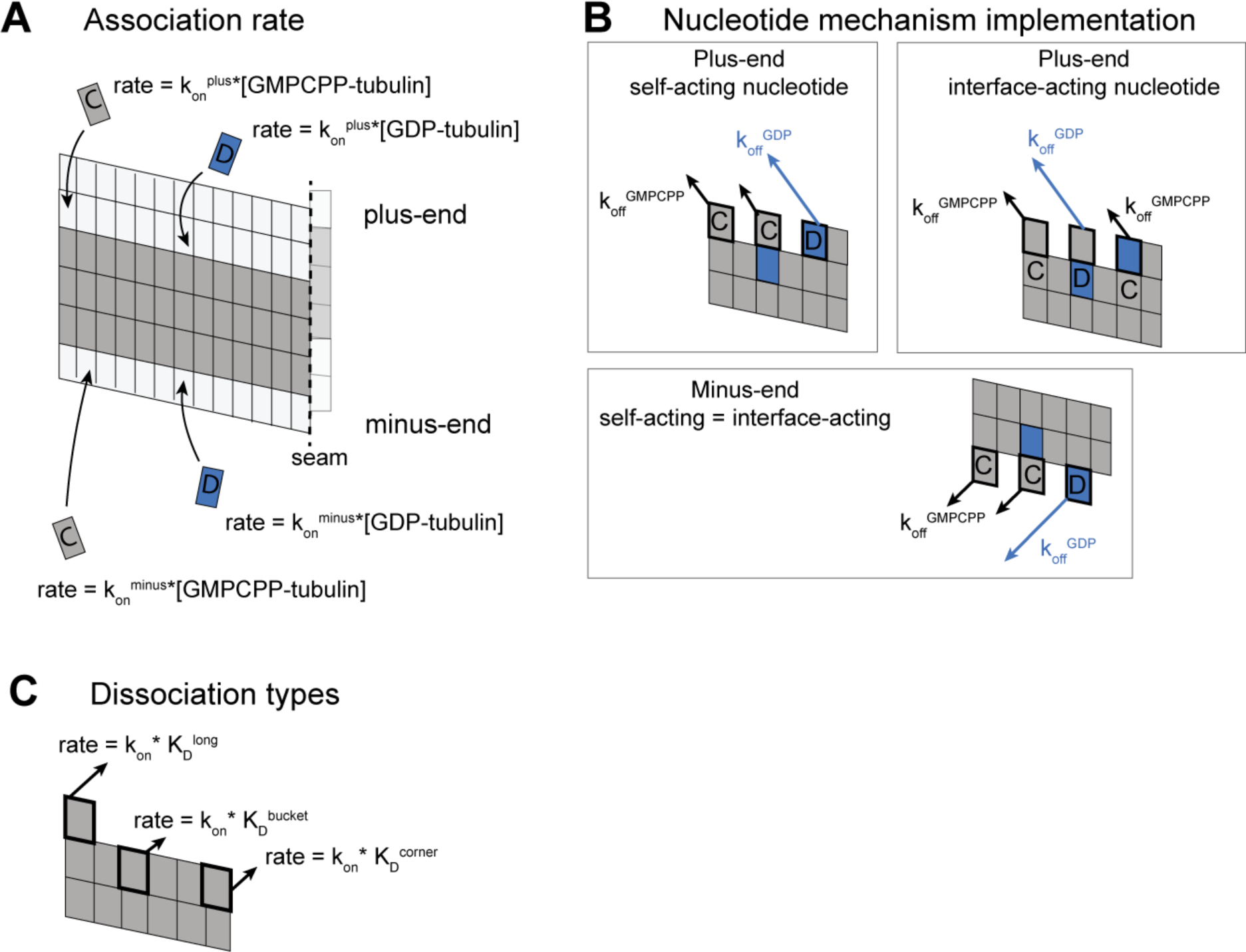
Implementation of plus- and minus-end models. **(A)** The microtubule lattice is represented as a two-dimensional grid; interactions between edge protofilaments generate the seam (dashed line) and mimic the cylindrical nature of the microtubule lattice. Grey boxes represent GMPCPP-tubulin, blue boxes represent GDP-tubulin, and white boxes represent empty (nucleotide free) positions. Arrows show how new subunits can associate at the plus-end or minus-end, respectively. Simulations begin with a microtubule seed, shown here as three rows of GMPCPP-tubulin. Subunit on-rates are determined by the on-rate constant (k_on_^plus^ or k_on_^minus^) and the concentration of tubulin. **(B)** Implementation of self-acting and interface-acting nucleotide mechanisms in plus-end simulations. Arrows indicate tubulin off-rates from the lattice, with black arrows denoting GMPCPP-tubulin off-rates and blue arrows denoting GDP-tubulin off-rates. **(C)** Tubulin dissociation rates from the lattice vary with the number of nearest neighbors. Plus-end simulations use the plus-end on-rate constant (k_on_^plus^), and minus-end simulations use the minus-end on-rate constant (k_on_^minus^).

**Figure 1 Supplement 2.**
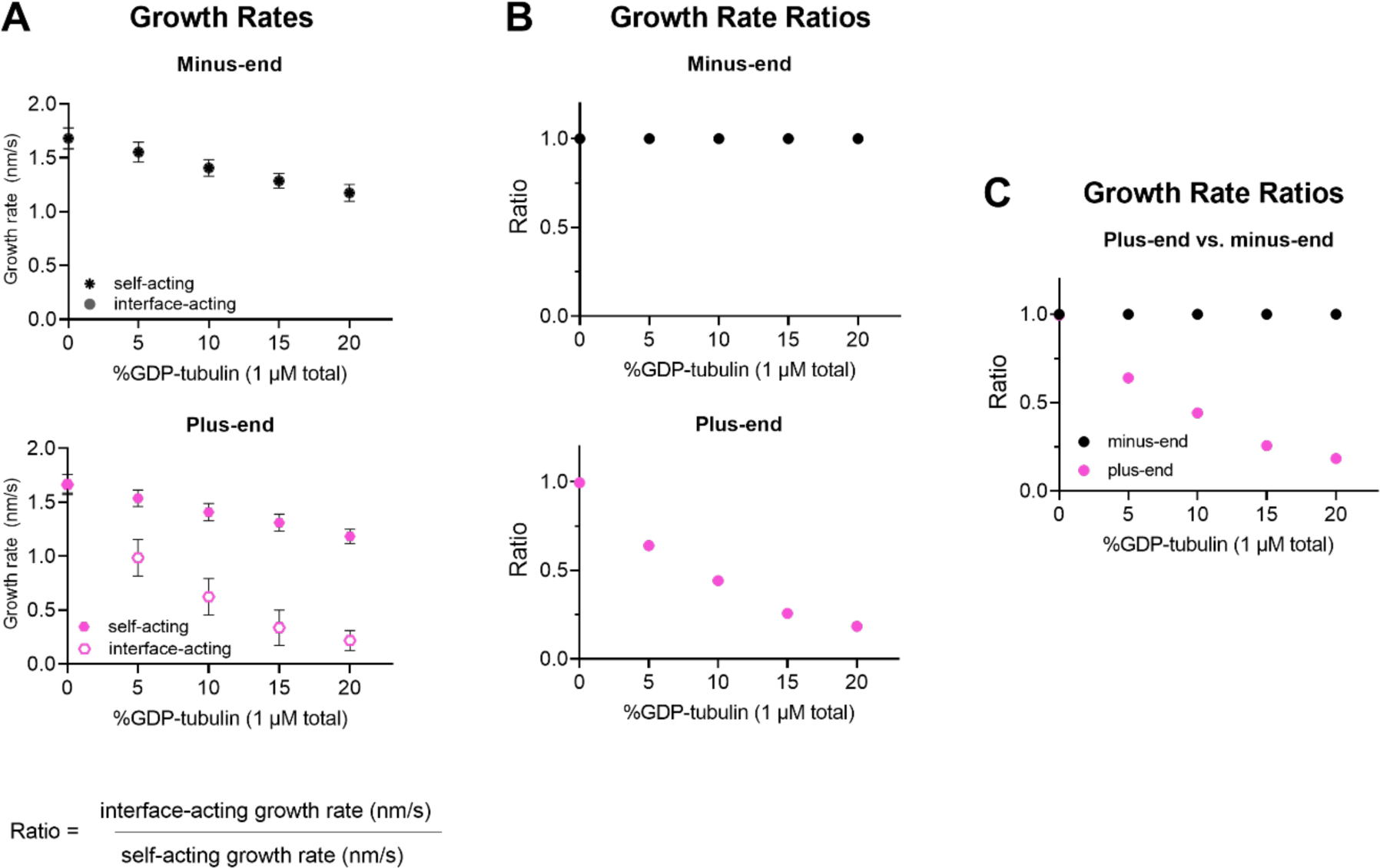
Using simulated growth rates to predict differences between interface- and self-acting nucleotide mechanisms at plus-end and minus-ends. **A)** Simulated minus-end and plus-end growth rates (nm/s) for interface-acting or self-acting nucleotide mechanisms using parameters shown in Figure 1. If growth rate markers are not visible, they are obscured by another marker. Error bars are standard deviation, with n = 50 independent simulations per concentration of GDP-tubulin. **B)** Growth rate ratios are defined as the growth rate for the interface-acting mechanism divided by the growth rate for the self-acting nucleotide mechanism, as a function of the GDP-tubulin concentration. A ratio of 1 indicates that no difference in growth rates is predicted for self- and interface-acting nucleotide mechanisms. **C)** Growth rate ratios of plus- and minus-ends from panel **B)** plotted together to emphasize how self- and interface-acting mechanisms predict increasingly different plus-end growth rates with increasing GDP-tubulin, whereas the two mechanisms predict similar minus-end growth rates across a range of GDP-tubulin concentrations.

**Figure 1 Supplement 3.**
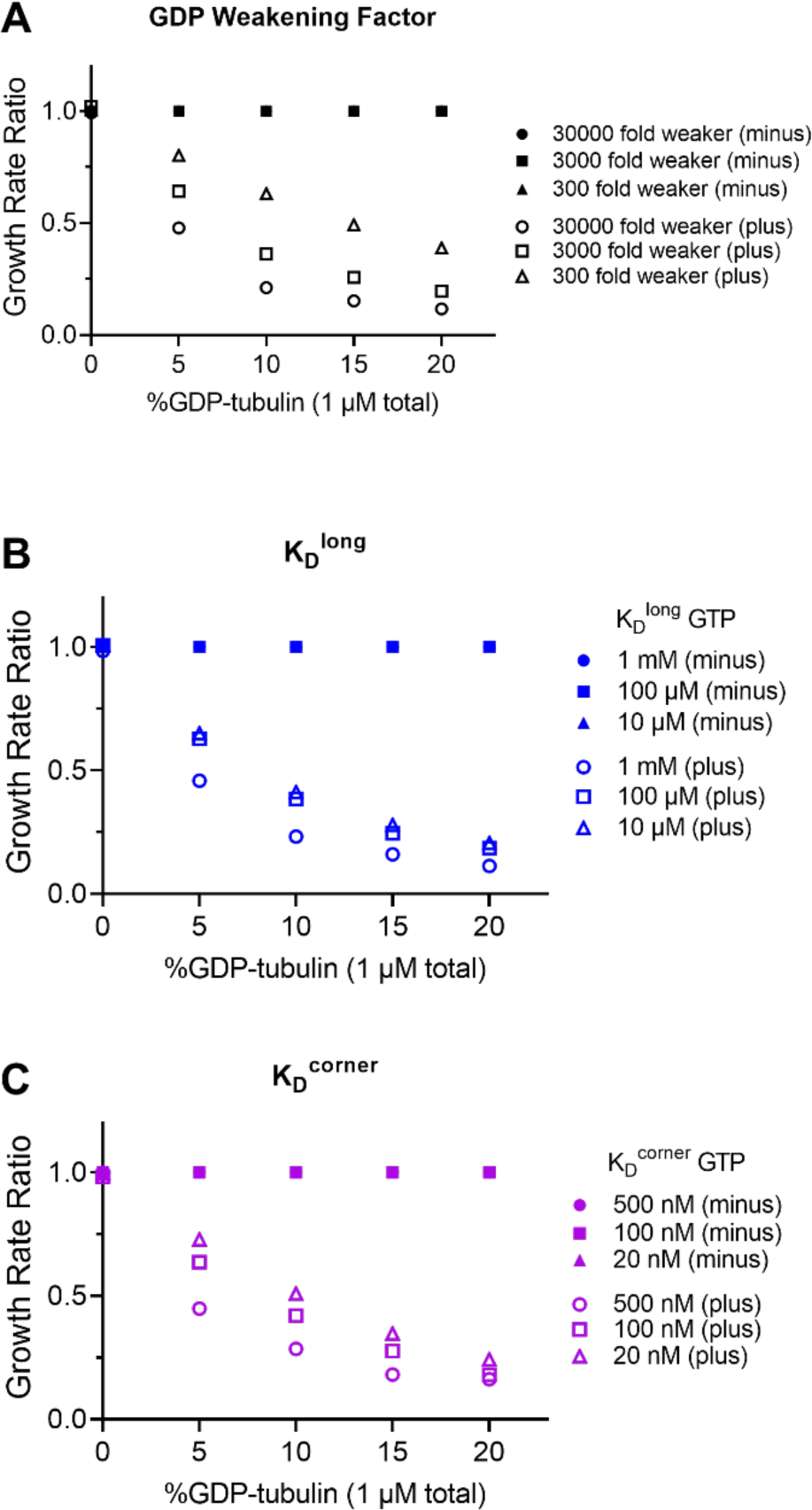
Predicted differences between self- and interface-acting mechanisms at the plus-end are robust to variation in simulation parameters. **A-C)** Ratios of simulated plus-end (open symbols) and minus-end (filled symbols) growth rates for interface- and self-acting nucleotide mechanisms. Growth rate ratios are calculated by dividing the interface-acting growth rate by the self-acting growth rate. A ratio of 1 means that no difference in growth rates is predicted. For each simulation parameter, a weaker and stronger choice (relative to the value used in Figure 1) was tested. The predicted difference between interface-acting and self-acting mechanisms persists, even for different choices of **A)** GDP weakening factor (100-fold range), **B)** longitudinal interaction (K_D_^long^, 100-fold range), and **C)** corner interaction (K_D_^corner^, 25-fold range). The original conditions used in Figure 1 are: GDP weakening factor of 3000, K_D_^long^ of 100 µM, and a K_D_^corner^ of 100 nM. The GDP weakening factor describes the fold change between the GDP- and GTP-type interactions. N = 50 simulations per concentration.

**Figure 3 – figure supplement 1.**
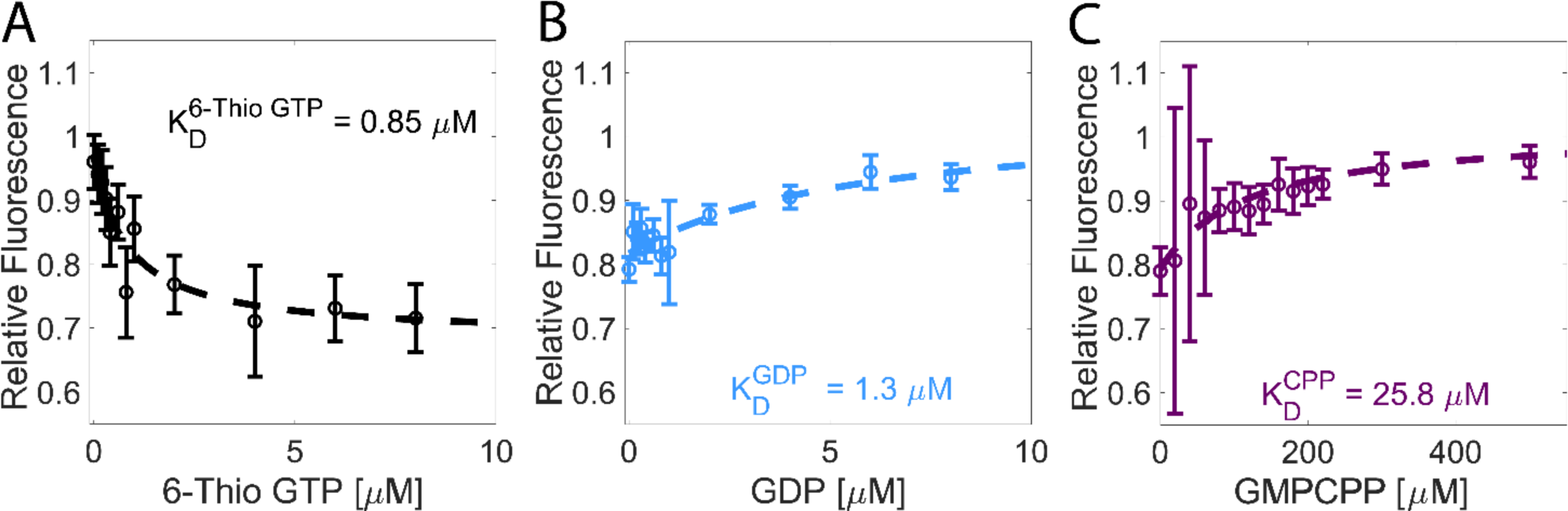
Tubulin has a higher affinity for GDP than for GMPCPP. **(A)** The affinity of tubulin for 6-Thio GTP measured by nucleotide-dependent quenching of the tubulin tryptophan fluorescence. Values are the tubulin fluorescence (0.2 µM tubulin) divided by the fluorescence of a signal-matched BSA sample to correct for the inner filter effect. Error bars are SEM for n=6 determinations for each sample, accounting for errors in the concentrations of tubulin and BSA, and in the buffer control. **(B)** Determination of tubulin affinity for GDP. Increasing concentrations of GDP were added to a solution of 0.2 µM tubulin in the presence of 3 µM 6-Thio GTP. GDP displaces 6-Thio GTP from the tubulin, causing unquenching of tryptophan fluorescence. Error bars denote SEM with n=6-11 determinations per point. **(C)** Determination of tubulin affinity for GMPCPP using an identical approach. Error bars are SEM with n=6 determinations per point.

**Figure 5 – figure supplement 1.**
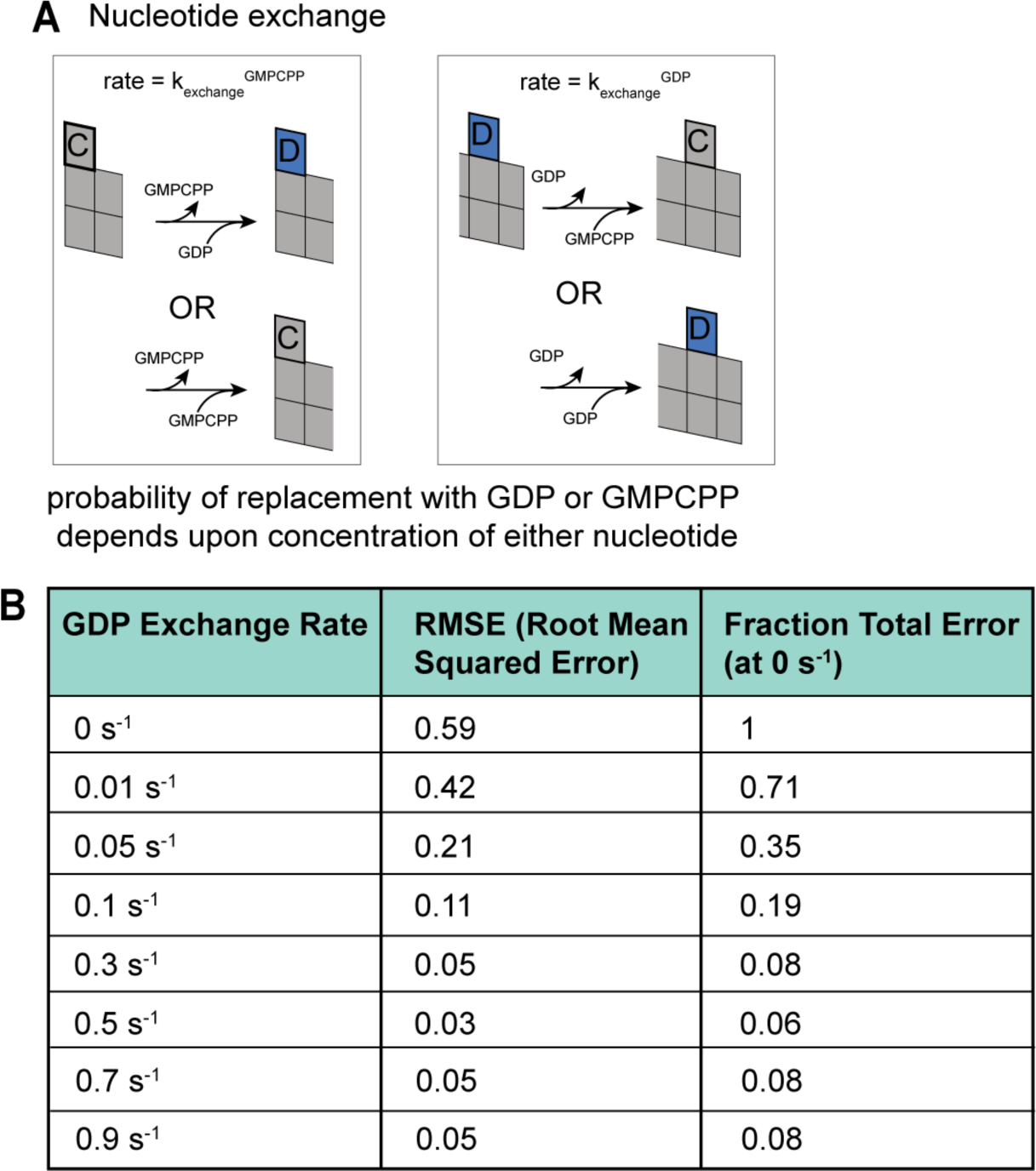
Implementation and analysis of nucleotide exchange. **(A)** Implementation of nucleotide exchange in simulations of plus-ended growth. Terminal exposed subunits can undergo exchange with a finite first-order rate, k_exchange_ (s^-1^). The probability of exchange to GDP or GMPCPP is determined by the relative concentration of each nucleotide. **(B)** Root-mean-squared error (RMSE) of predicted growth rates vs. experimental growth rates for a series of exchange rates, across the tested range of GDP- and GMPCPP-tubulin mixtures. Fraction total error is defined as the relative error compared to the error when exchange rate is 0 s^-1^.

## Notes

### Competing Interest Statement

The authors have declared no competing interest.

### Summary of Updates

This version of the manuscript has been revised to address comments from reviewers.

## Bibliography

Akhmanova, A., & Kapitein, L. C. (2022). Mechanisms of microtubule organization in differentiated animal cells. Nat Rev Mol Cell Biol, 23(8), 541–558. 10.1038/s41580-022-00473-y

Aldaz, H., Rice, L. M., Stearns, T., & Agard, D. A. (2005). Insights into microtubule nucleation from the crystal structure of human gamma-tubulin. Nature, 435(7041), 523–527. 10.1038/nature03586

Alushin, G. M., Lander, G. C., Kellogg, E. H., Zhang, R., Baker, D., & Nogales, E. (2014). High-resolution microtubule structures reveal the structural transitions in alphabeta-tubulin upon GTP hydrolysis. Cell, 157(5), 1117–1129. 10.1016/j.cell.2014.03.053

Amayed, P., Carlier, M. F., & Pantaloni, D. (2000). Stathmin slows down guanosine diphosphate dissociation from tubulin in a phosphorylation-controlled fashion. Biochemistry, 39(40), 12295–12302. 10.1021/bi000279w

Ayaz, P., Munyoki, S., Geyer, E. A., Piedra, F. A., Vu, E. S., Bromberg, R., Otwinowski, Z., Grishin, N. V., Brautigam, C. A., & Rice, L. M. (2014). A tethered delivery mechanism explains the catalytic action of a microtubule polymerase. Elife, 3, e03069. 10.7554/eLife.03069

Ayaz, P., Ye, X., Huddleston, P., Brautigam, C. A., & Rice, L. M. (2012). A TOG:alphabeta-tubulin complex structure reveals conformation-based mechanisms for a microtubule polymerase. Science, 337(6096), 857–860. 10.1126/science.1221698

Banerjee, S., Bartesaghi, A., Merk, A., Rao, P., Bulfer, S. L., Yan, Y., Green, N., Mroczkowski, B., Neitz, R. J., Wipf, P., Falconieri, V., Deshaies, R. J., Milne, J. L., Huryn, D., Arkin, M., & Subramaniam, S. (2016). 2.3 A resolution cryo-EM structure of human p97 and mechanism of allosteric inhibition. Science, 351(6275), 871–875. 10.1126/science.aad7974

Barlan, K., & Gelfand, V. I. (2017). Microtubule-Based Transport and the Distribution, Tethering, and Organization of Organelles. Cold Spring Harb Perspect Biol, 9(5). 10.1101/cshperspect.a025817

Bowne-Anderson, H., Zanic, M., Kauer, M., & Howard, J. (2013). Microtubule dynamic instability: a new model with coupled GTP hydrolysis and multistep catastrophe. Bioessays, 35(5), 452–461. 10.1002/bies.201200131

Brouhard, G., & Sept, D. (2012). Microtubules: sizing up the GTP cap. Curr Biol, 22(18), R802–803. 10.1016/j.cub.2012.07.050

Brouhard, G. J. (2015). Dynamic instability 30 years later: complexities in microtubule growth and catastrophe. Mol Biol Cell, 26(7), 1207–1210. 10.1091/mbc.E13-10-0594

Brouhard, G. J., & Rice, L. M. (2018). Microtubule dynamics: an interplay of biochemistry and mechanics. Nat Rev Mol Cell Biol, 19(7), 451–463. 10.1038/s41580-018-0009-y

Brun, L., Rupp, B., Ward, J. J., & Nedelec, F. (2009). A theory of microtubule catastrophes and their regulation. Proc Natl Acad Sci U S A, 106(50), 21173–21178. 10.1073/pnas.0910774106

Brunger, A. T., & DeLaBarre, B. (2003). NSF and p97/VCP: similar at first, different at last. FEBS Lett, 555(1), 126–133. 10.1016/s0014-5793(03)01107-4

Buey, R. M., Diaz, J. F., & Andreu, J. M. (2006). The nucleotide switch of tubulin and microtubule assembly: a polymerization-driven structural change. Biochemistry, 45(19), 5933–5938. 10.1021/bi060334m

Caplow, M., & Shanks, J. (1996). Evidence that a single monolayer tubulin-GTP cap is both necessary and sufficient to stabilize microtubules. Mol Biol Cell, 7(4), 663–675. 10.1091/mbc.7.4.663

Carlier, M. F., & Pantaloni, D. (1978). Kinetic analysis of cooperativity in tubulin polymerization in the presence of guanosine di- or triphosphate nucleotides. Biochemistry, 17(10), 1908–1915. 10.1021/bi00603a017

Chakrabarti, G., Mejillano, M. R., Park, Y. H., Vander Velde, D. G., & Himes, R. H. (2000). Nucleoside triphosphate specificity of tubulin. Biochemistry, 39(33), 10269–10274. 10.1021/bi000966n

Chen, Y., & Hill, T. L. (1983). Use of Monte Carlo calculations in the study of microtubule subunit kinetics. Proc Natl Acad Sci U S A, 80(24), 7520–7523. 10.1073/pnas.80.24.7520

Chen, Y. D., & Hill, T. L. (1985). Monte Carlo study of the GTP cap in a five-start helix model of a microtubule. Proc Natl Acad Sci U S A, 82(4), 1131–1135. 10.1073/pnas.82.4.1131

Cheng, Y., & Prusoff, W. H. (1973). Relationship between the inhibition constant (K1) and the concentration of inhibitor which causes 50 per cent inhibition (I50) of an enzymatic reaction. Biochem Pharmacol, 22(23), 3099–3108. 10.1016/0006-2952(73)90196-2

Cleary, J. M., & Hancock, W. O. (2021). Molecular mechanisms underlying microtubule growth dynamics. Curr Biol, 31(10), R560–R573. 10.1016/j.cub.2021.02.035

Cleary, J. M., Kim, T., Cook, A. S. I., McCormick, L. A., Hancock, W. O., & Rice, L. M. (2022). Measurements and simulations of microtubule growth imply strong longitudinal interactions and reveal a role for GDP on the elongating end. Elife, 11. 10.7554/eLife.75931

Coombes, C. E., Yamamoto, A., Kenzie, M. R., Odde, D. J., & Gardner, M. K. (2013). Evolving tip structures can explain age-dependent microtubule catastrophe. Curr Biol, 23(14), 1342–1348. 10.1016/j.cub.2013.05.059

Correia, J. J., Baty, L. T., & Williams, R. C., Jr. (1987). Mg2+ dependence of guanine nucleotide binding to tubulin. J Biol Chem, 262(36), 17278–17284. https://www.ncbi.nlm.nih.gov/pubmed/2826416

Cross, R. A. (2019). Microtubule lattice plasticity. Curr Opin Cell Biol, 56, 88–93. 10.1016/j.ceb.2018.10.004

Davies, J. M., Tsuruta, H., May, A. P., & Weis, W. I. (2005). Conformational changes of p97 during nucleotide hydrolysis determined by small-angle X-Ray scattering. Structure, 13(2), 183–195. 10.1016/j.str.2004.11.014

Drechsel, D. N., & Kirschner, M. W. (1994). The minimum GTP cap required to stabilize microtubules. Curr Biol, 4(12), 1053–1061. 10.1016/s0960-9822(00)00243-8

Duellberg, C., Cade, N. I., & Surrey, T. (2016). Microtubule aging probed by microfluidics-assisted tubulin washout. Mol Biol Cell, 27(22), 3563–3573. 10.1091/mbc.E16-07-0548

Erzberger, J. P., & Berger, J. M. (2006). Evolutionary relationships and structural mechanisms of AAA+ proteins. Annu Rev Biophys Biomol Struct, 35, 93–114. 10.1146/annurev.biophys.35.040405.101933

Farmer, V., Arpag, G., Hall, S. L., & Zanic, M. (2021). XMAP215 promotes microtubule catastrophe by disrupting the growing microtubule end. J Cell Biol, 220(10). 10.1083/jcb.202012144

Farmer, V. J., & Zanic, M. (2023). Beyond the GTP-cap: Elucidating the molecular mechanisms of microtubule catastrophe. Bioessays, 45(1), e2200081. 10.1002/bies.202200081

Fishback, J. L., & Yarbrough, L. R. (1984). Interaction of 6-mercapto-GTP with bovine brain tubulin. Equilibrium aspects. J Biol Chem, 259(3), 1968–1973. https://www.ncbi.nlm.nih.gov/pubmed/6693440

Gudimchuk, N. B., & McIntosh, J. R. (2021). Regulation of microtubule dynamics, mechanics and function through the growing tip. Nat Rev Mol Cell Biol, 22(12), 777–795. 10.1038/s41580-021-00399-x

Hamel, E., Batra, J. K., & Lin, C. M. (1986). Direct incorporation of guanosine 5’-diphosphate into microtubules without guanosine 5’-triphosphate hydrolysis. Biochemistry, 25(22), 7054–7062. 10.1021/bi00370a045

Howard, J., & Hyman, A. A. (2009). Growth, fluctuation and switching at microtubule plus ends. Nat Rev Mol Cell Biol, 10(8), 569–574. 10.1038/nrm2713

Howard, W. D., & Timasheff, S. N. (1986). GDP state of tubulin: stabilization of double rings. Biochemistry, 25(25), 8292–8300. 10.1021/bi00373a025

Hyman, A. A., Salser, S., Drechsel, D. N., Unwin, N., & Mitchison, T. J. (1992). Role of GTP hydrolysis in microtubule dynamics: information from a slowly hydrolyzable analogue, GMPCPP. Mol Biol Cell, 3(10), 1155–1167. 10.1091/mbc.3.10.1155

Kim, T., & Rice, L. M. (2019). Long-range, through-lattice coupling improves predictions of microtubule catastrophe. Mol Biol Cell, 30(12), 1451–1462. 10.1091/mbc.E18-10-0641

LaFrance, B. J., Roostalu, J., Henkin, G., Greber, B. J., Zhang, R., Normanno, D., McCollum, C. O., Surrey, T., & Nogales, E. (2022). Structural transitions in the GTP cap visualized by cryo-electron microscopy of catalytically inactive microtubules. Proc Natl Acad Sci U S A, 119(2). 10.1073/pnas.2114994119

Lawrence, E., Chatterjee, S., & Zanic, M. (2022). CLASPs stabilize the intermediate state between microtubule growth and catastrophe. bioRxiv, 2022.2012.2003.518990. 10.1101/2022.12.03.518990

Luo, W., Demidov, V., Shen, Q., Girao, H., Chakraborty, M., Maiorov, A., Ataullakhanov, F. I., Lin, C., Maiato, H., & Grishchuk, E. L. (2023). CLASP2 recognizes tubulins exposed at the microtubule plus-end in a nucleotide state-sensitive manner. Sci Adv, 9(1), eabq5404. 10.1126/sciadv.abq5404

Mahserejian, S. M., Scripture, J. P., Mauro, A. J., Lawrence, E. J., Jonasson, E. M., Murray, K. S., Li, J., Gardner, M., Alber, M., Zanic, M., & Goodson, H. V. (2022). Quantification of microtubule stutters: dynamic instability behaviors that are strongly associated with catastrophe. Mol Biol Cell, 33(3), ar22. 10.1091/mbc.E20-06-0348

Manka, S. W., & Moores, C. A. (2018). The role of tubulin-tubulin lattice contacts in the mechanism of microtubule dynamic instability. Nat Struct Mol Biol, 25(7), 607–615. 10.1038/s41594-018-0087-8

Margolin, G., Gregoretti, I. V., Cickovski, T. M., Li, C., Shi, W., Alber, M. S., & Goodson, H. V. (2012). The mechanisms of microtubule catastrophe and rescue: implications from analysis of a dimer-scale computational model. Mol Biol Cell, 23(4), 642–656. 10.1091/mbc.E11-08-0688

Maurer, S. P., Fourniol, F. J., Bohner, G., Moores, C. A., & Surrey, T. (2012). EBs recognize a nucleotide-dependent structural cap at growing microtubule ends. Cell, 149(2), 371–382. 10.1016/j.cell.2012.02.049

Mejillano, M. R., & Himes, R. H. (1991). Binding of guanine nucleotides and Mg2+ to tubulin with a nucleotide-depleted exchangeable site. Arch Biochem Biophys, 291(2), 356–362. 10.1016/0003-9861(91)90146-a

Melki, R., Carlier, M. F., Pantaloni, D., & Timasheff, S. N. (1989). Cold depolymerization of microtubules to double rings: geometric stabilization of assemblies. Biochemistry, 28(23), 9143–9152. 10.1021/bi00449a028

Mickolajczyk, K. J., Geyer, E. A., Kim, T., Rice, L. M., & Hancock, W. O. (2019). Direct observation of individual tubulin dimers binding to growing microtubules. Proc Natl Acad Sci U S A, 116(15), 7314–7322. 10.1073/pnas.1815823116

Mitchison, T., & Kirschner, M. (1984). Dynamic instability of microtubule growth. Nature, 312(5991), 237–242. 10.1038/312237a0

Mitchison, T. J. (1993). Localization of an exchangeable GTP binding site at the plus end of microtubules. Science, 261(5124), 1044–1047. 10.1126/science.8102497

Molodtsov, M. I., Ermakova, E. A., Shnol, E. E., Grishchuk, E. L., McIntosh, J. R., & Ataullakhanov, F. I. (2005). A molecular-mechanical model of the microtubule. Biophys J, 88(5), 3167–3179. 10.1529/biophysj.104.051789

Monasterio, O., & Timasheff, S. N. (1987). Inhibition of tubulin self-assembly and tubulin-colchicine GTPase activity by guanosine 5’-(gamma-fluorotriphosphate). Biochemistry, 26(19), 6091–6099. 10.1021/bi00393a022

Nawrotek, A., Knossow, M., & Gigant, B. (2011). The determinants that govern microtubule assembly from the atomic structure of GTP-tubulin. J Mol Biol, 412(1), 35–42. 10.1016/j.jmb.2011.07.029

Nicholson, W. V., Lee, M., Downing, K. H., & Nogales, E. (1999). Cryo-electron microscopy of GDP-tubulin rings. Cell Biochem Biophys, 31(2), 175–183. 10.1007/BF02738171

Pecqueur, L., Duellberg, C., Dreier, B., Jiang, Q., Wang, C., Pluckthun, A., Surrey, T., Gigant, B., & Knossow, M. (2012). A designed ankyrin repeat protein selected to bind to tubulin caps the microtubule plus end. Proc Natl Acad Sci U S A, 109(30), 12011–12016. 10.1073/pnas.1204129109

Piedra, F. A., Kim, T., Garza, E. S., Geyer, E. A., Burns, A., Ye, X., & Rice, L. M. (2016). GDP-to-GTP exchange on the microtubule end can contribute to the frequency of catastrophe. Mol Biol Cell, 27(22), 3515–3525. 10.1091/mbc.E16-03-0199

Prosser, S. L., & Pelletier, L. (2017). Mitotic spindle assembly in animal cells: a fine balancing act. Nat Rev Mol Cell Biol, 18(3), 187–201. 10.1038/nrm.2016.162

Rice, L. M., Montabana, E. A., & Agard, D. A. (2008). The lattice as allosteric effector: structural studies of alphabeta- and gamma-tubulin clarify the role of GTP in microtubule assembly. Proc Natl Acad Sci U S A, 105(14), 5378–5383. 10.1073/pnas.0801155105

Roostalu, J., Thomas, C., Cade, N. I., Kunzelmann, S., Taylor, I. A., & Surrey, T. (2020). The speed of GTP hydrolysis determines GTP cap size and controls microtubule stability. Elife, 9. 10.7554/eLife.51992

Roth, D., Fitton, B. P., Chmel, N. P., Wasiluk, N., & Straube, A. (2018). Spatial positioning of EB family proteins at microtubule tips involves distinct nucleotide-dependent binding properties. J Cell Sci, 132(4). 10.1242/jcs.219550

Ruhnow, F., Zwicker, D., & Diez, S. (2011). Tracking single particles and elongated filaments with nanometer precision. Biophys J, 100(11), 2820–2828. 10.1016/j.bpj.2011.04.023

Schindelin, J., Rueden, C. T., Hiner, M. C., & Eliceiri, K. W. (2015). The ImageJ ecosystem: An open platform for biomedical image analysis. Mol Reprod Dev, 82(7-8), 518–529. 10.1002/mrd.22489

Schmidt, M., & Kierfeld, J. (2021). Chemomechanical Simulation of Microtubule Dynamics With Explicit Lateral Bond Dynamics [Original Research]. Frontiers in Physics, 9. 10.3389/fphy.2021.673875

Seetapun, D., Castle, B. T., McIntyre, A. J., Tran, P. T., & Odde, D. J. (2012). Estimating the microtubule GTP cap size in vivo. Curr Biol, 22(18), 1681–1687. 10.1016/j.cub.2012.06.068

Shearwin, K. E., Perez-Ramirez, B., & Timasheff, S. N. (1994). Linkages between the dissociation of alpha beta tubulin into subunits and ligand binding: the ground state of tubulin is the GDP conformation. Biochemistry, 33(4), 885–893. 10.1021/bi00170a006

Stewman, S. F., Tsui, K. K., & Ma, A. (2020). Dynamic Instability from Non-equilibrium Structural Transitions on the Energy Landscape of Microtubule. Cell Syst, 11(6), 608–624 e609. 10.1016/j.cels.2020.09.008

Strothman, C., Farmer, V., Arpag, G., Rodgers, N., Podolski, M., Norris, S., Ohi, R., & Zanic, M. (2019). Microtubule minus-end stability is dictated by the tubulin off-rate. J Cell Biol, 218(9), 2841–2853. 10.1083/jcb.201905019

Tanaka-Takiguchi, Y., Itoh, T. J., & Hotani, H. (1998). Visualization of the GDP-dependent switching in the growth polarity of microtubules. J Mol Biol, 280(3), 365–373. 10.1006/jmbi.1998.1877

Tran, P. T., Joshi, P., & Salmon, E. D. (1997). How tubulin subunits are lost from the shortening ends of microtubules. J Struct Biol, 118(2), 107–118. 10.1006/jsbi.1997.3844

Uppalapati, M., Huang, Y.-M., Shastry, S., Jackson, T. N., & Hancock, W. O. (2009). Microtubule motors in microfluidics. Methods in bioengineering: microfabrication and microfluidics, 311–336.

Valiron, O., Arnal, I., Caudron, N., & Job, D. (2010). GDP-tubulin incorporation into growing microtubules modulates polymer stability. J Biol Chem, 285(23), 17507–17513. 10.1074/jbc.M109.099515

VanBuren, V., Cassimeris, L., & Odde, D. J. (2005). Mechanochemical model of microtubule structure and self-assembly kinetics. Biophys J, 89(5), 2911–2926. 10.1529/biophysj.105.060913

VanBuren, V., Odde, D. J., & Cassimeris, L. (2002). Estimates of lateral and longitudinal bond energies within the microtubule lattice. Proc Natl Acad Sci U S A, 99(9), 6035–6040. 10.1073/pnas.092504999

Vandecandelaere, A., Martin, S. R., & Bayley, P. M. (1995). Regulation of microtubule dynamic instability by tubulin-GDP. Biochemistry, 34(4), 1332–1343. 10.1021/bi00004a028

Walker, R. A., O’Brien, E. T., Pryer, N. K., Soboeiro, M. F., Voter, W. A., Erickson, H. P., & Salmon, E. D. (1988). Dynamic instability of individual microtubules analyzed by video light microscopy: rate constants and transition frequencies. J Cell Biol, 107(4), 1437–1448. 10.1083/jcb.107.4.1437

Wang, H. W., & Nogales, E. (2005). Nucleotide-dependent bending flexibility of tubulin regulates microtubule assembly. Nature, 435(7044), 911–915. 10.1038/nature03606

Yarbrough, L. R., & Fishback, J. L. (1985). Kinetics of interaction of 2-amino-6-mercapto-9-beta-ribofuranosylpurine 5’-triphosphate with bovine brain tubulin. Biochemistry, 24(7), 1708–1714. 10.1021/bi00328a021

Zakharov, P., Gudimchuk, N., Voevodin, V., Tikhonravov, A., Ataullakhanov, F. I., & Grishchuk, E. L. (2015). Molecular and Mechanical Causes of Microtubule Catastrophe and Aging. Biophys J, 109(12), 2574–2591. 10.1016/j.bpj.2015.10.048

Zanic, M., Widlund, P. O., Hyman, A. A., & Howard, J. (2013). Synergy between XMAP215 and EB1 increases microtubule growth rates to physiological levels. Nat Cell Biol, 15(6), 688–693. 10.1038/ncb2744

Zeeberg, B., & Caplow, M. (1979). Determination of free and bound microtubular protein and guanine nucleotide under equilibrium conditions. Biochemistry, 18(18), 3880–3886. 10.1021/bi00585a007

Zhang, R., Alushin, G. M., Brown, A., & Nogales, E. (2015). Mechanistic Origin of Microtubule Dynamic Instability and Its Modulation by EB Proteins. Cell, 162(4), 849–859. 10.1016/j.cell.2015.07.012

